# Genetic exchange in *Leishmania* is facilitated by IgM natural antibodies

**DOI:** 10.1101/2022.06.09.495557

**Authors:** Tiago D. Serafim, Eva Iniguez, Ana Beatriz F. Barletta, Johannes S.P. Doehl, Mara Short, Justin Lack, Pedro Cecilio, Vinod Nair, Maria Disotuar, Timothy Wilson, Iliano V. Coutinho-Abreu, Fabiano Oliveira, Claudio Meneses, Carolina Barillas-Mury, John Andersen, José M.C. Ribeiro, Stephen M. Beverley, Shaden Kamhawi, Jesus G. Valenzuela

**Affiliations:** Vector Molecular Biology Section, Laboratory of Malaria and Vector Research, National Institute of Allergy and Infectious Diseases, National Institutes of Health, Rockville, MD, USA; Mosquito Immunity and Vector Competence Section, Laboratory of Malaria and Vector Research, National Institute of Allergy and Infectious Diseases, National Institutes of Health, Rockville, MD, USA; NIAID Collaborative Bioinformatics Resource, National Institute of Allergy and Infectious Diseases, National Institutes of Health, Bethesda, MD, USA; Vector Biology Section, Laboratory of Malaria and Vector Research, National Institute of Allergy and Infectious Diseases, National Institutes of Health, Rockville, MD, USA; Rocky Mountain Laboratories, National Institute of Allergy and Infectious Diseases, National Institutes of Health, Hamilton, MT, USA; Department of Molecular Microbiology, School of Medicine, Washington University, St. Louis, MO, USA

## Abstract

Host factors mediating *Leishmania* genetic exchange are not well defined. Here, we demonstrate that IgM antibodies, but not IgG or IgA, facilitate parasite genetic hybridization in vitro and in vivo. IgM induces the gradual and transient formation of a structured parasite clump in a process essential for *L. major* and *L. tropica* hybridization in vitro. Parasite hybrids and 3-nucleated parasites were observed inside this structure, named the *Leishmania* mating clump. IgM was also required for or significantly increased *Leishmania* hybrid formation in vivo. At minimum, we observed a 12-fold increase in the proportion of hybrids recovered from sand flies provided a second blood meal containing IgM compared to controls. Notably, genetic backcross events in sand flies were only observed in the presence of IgM, and were reproducibly recovered, reinforcing the relevance of IgM for *Leishmania* genetic exchange in vivo. The in vitro and in vivo *Leishmania* crosses from these studies resulted in full genome hybrids. *Leishmania* co-option of a host antibody to facilitate mating in the insect vector establishes a new paradigm of parasite-host-vector coevolution that promotes parasite diversity and fitness through genetic exchange.

## Main text

*Leishmania* parasites are vector-borne pathogens transmitted to mammalian hosts by phlebotomine sand flies^1^. Associated with poverty, leishmaniasis is prevalent in developing countries and results in self-limiting, severely disfiguring, or fatal diseases that cause substantial morbidity and mortality worldwide^2^. More than a billion people are at risk of infection, millions are asymptomatic carriers, and the annual incidence of fatal visceral and cutaneous leishmaniases are estimated at 100,000 to 1 million cases, respectively^3^. These statistics place leishmaniasis at the forefront of parasitic diseases of public health importance.

*Leishmania* parasites are considered mostly asexual, reproducing via clonal propagation^4–6^, though the presence of natural hybrids and demonstration of genetic exchange in the sand fly vector^7–10^ indicate that genetic exchange is more likely ‘facultative’ as seen in other eukaryotic microbes^11^. Genetic exchange is a powerful evolutionary route enabling the development of pathogenicity in microbes and offers an experimental tool for dissecting complex phenotypes underlying virulence^12^.

Host and parasite factors mediating this process are not well understood. In earlier studies, passage in sand flies was required to undertake this experimentally^10,13,14^, but recently, hybrid formation was observed in culture for a few species including *L. tropica*, albeit following exposure to harsh conditions^10,15,16^. Nevertheless, in vitro hybrid generation is yet to be accomplished for most *Leishmania* species, including *L. major*^10,15,16^. Importantly, genetic exchange or the generation of fertile hybrids through backcrosses has only been reported once^8^.

In this study, we focused on elucidating host factors that mediate *Leishmania* genetic exchange, asking whether host blood ingested by sand flies provides some element(s) absent in culture media that promotes fusion and the formation of hybrid parasites. By adding freshly isolated dog plasma to culture media we succeeded in generating in vitro crosses of two genetically labeled *L. major* strains, WR-SSU-HYG and FV1-FKP40-BSD (Fig. 1a). Hybrid generation was only observed when accompanied by the formation of a structured spherical ‘clump’ composed of live aggregating parasites (Fig. 1b, top panel; Supplementary Videos 1-3). In standard culture media, in the absence of dog plasma, the parasite clump did not form (Fig. 1b, bottom panel; Supplementary Videos 6,7) and hybrids were not recovered. The parasite clump forms gradually and is maintained for 24-36h (Supplementary Video 4) before it starts to dissociate releasing viable parasites.

**Fig 1.**
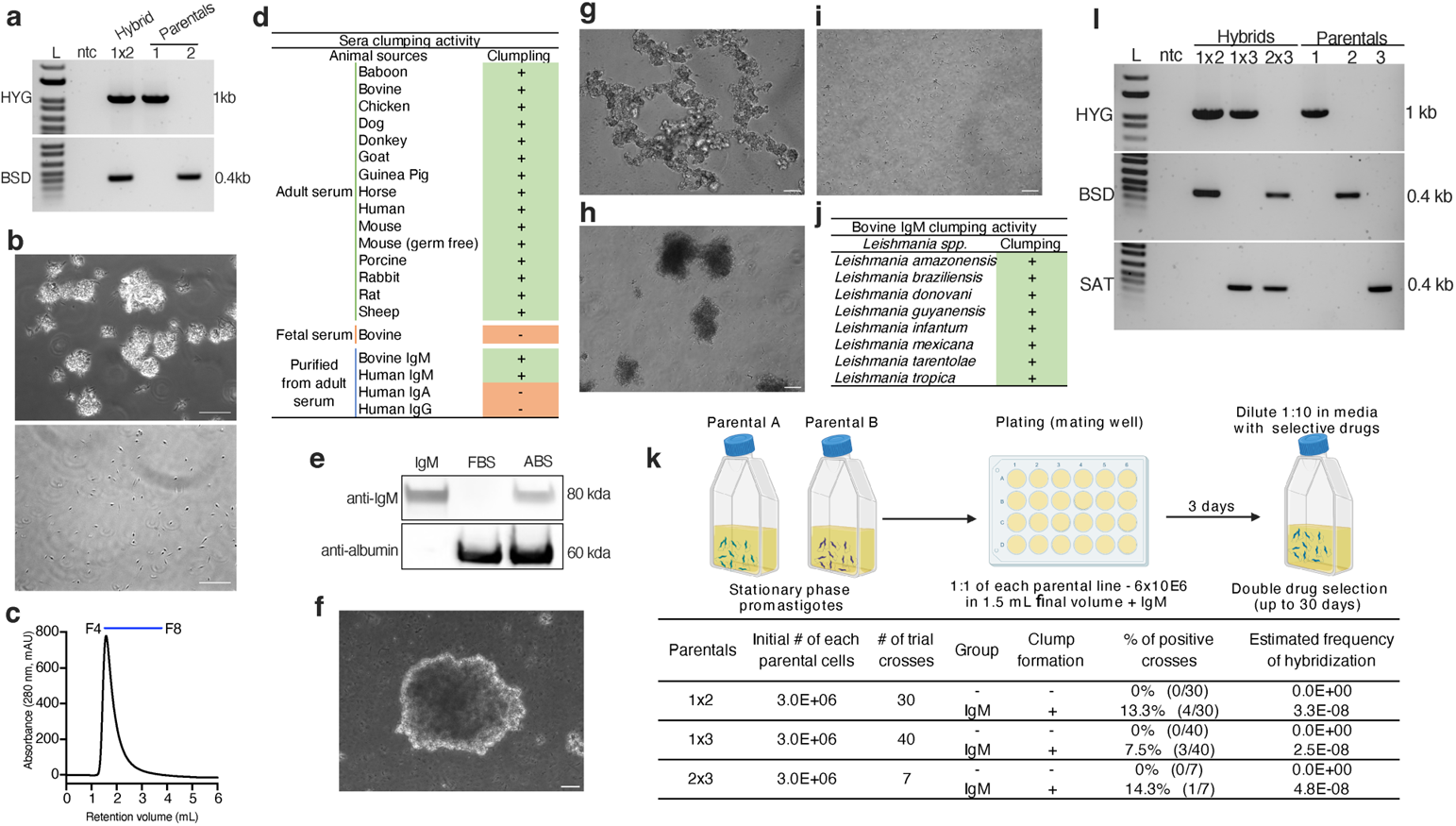
IgM facilitates *Leishmania* parasites clumping which promotes hybrid formation in vitro. (**a**) *Leishmania* hybrids genotyping by PCR targeting parental selectable drug markers HYG (Hygromycin) and BSD (Blasticidin). (**b**) Phase contrast image of *L. major* in culture media containing 20% fetal bovine serum (FBS) without (bottom panel) or with 5% adult inactivated dog serum (top panel). Images taken 30 min after cells were seeded in fresh media. Scale bars = 20 µm. (**c**) Human IgM eluted from a HiTrap® IgM column. *Leishmania* clumping activity was observed in fractions F4-F8 corresponding to elution of IgM. **(d)** Presence (+) or absence (−) of *L. major* clumping activity in inactivated adult sera from different vertebrate species, FBS, or immunoglobulins (IgM, IgG, IgA) purified from adult bovine or human sera. (**e**) Western blot to detect IgM in FBS and adult bovine serum (ABS). Serum purified bovine IgM, positive control; serum albumin, loading control. (**f**) Phase contrast image of *L. major* in culture media containing 20% FBS with bovine IgM at 50 µg/mL after 24h and (**g**) with peanut agglutinin (PNA) at 50 µg/mL after 24h. Scale bars = 50 µm. (**h**) Phase contrast image of *L. major* in culture media containing 20% FBS with bovine IgM (pentameric form) at 50 µg/mL after 24h or (**i**) with bovine IgM (monomeric form) at 50 µg/mL after 24h. (**j**) Clumping activity of various *Leishmania* species in the presence of bovine IgM at 50 µg/mL. (**k**) (top) Workflow for in vitro crossing of *L. major* parental lines. Mating wells were established with 50 µg/mL IgM in complete Schneider’s media. Double drug resistant hybrid parasites were cloned before genotyping. (bottom) Summary of data from in vitro crossing of parental lines. 1×2 (n=30), 1×3 (n=40), 2×3 (n=7). (**l**) *Leishmania major* hybrids genotyping by PCR targeting parental selectable drug markers HYG (Hygromycin), BSD (Blasticidin) and SAT (Nourseothricin). Parental 1, WR-SSU-HYG; Parental 2, FVI-FKP40-BSD; parental 3, FVI-FTL-SAT; ntc, no template control; L, 1kb plus ladder (Invitrogen).

Purification and fractionation of dog plasma using a 2 step-HPLC followed by MS-MS and in vitro incubation of parasites with the HPLC fractions revealed that IgM was the molecule responsible for the formation of the parasite clump (Supplementary Fig. 1). Parasite clumping was retained in HPLC-eluted fractions of a commercially available IgM (Fig. 1c) but was absent with a commercial human IgA or IgG (Fig. 1d), emphasizing the unique role of IgM in parasite clumping. Importantly, fetal bovine serum (FBS) does not induce clumping (Fig. 1d) and it does not contain IgM (Fig. 1e). Pertinently, the process of IgM-induced clump formation (Fig. 1f, Supplementary Fig. 2a) is distinct from other aggregation phenomena such as with peanut agglutinin (PNA), where the parasites agglutinate immediately and irreversibly, forming a disorganized flat mesh-like structure of immotile parasites (Fig. 1g; Supplementary Fig. 2b; Supplementary Video 5). The composition and temporal formation and dissociation of the *Leishmania* parasite clump is also different from other reported structures, including the *Leishmania* rosettes^17^. Furthermore, pentameric IgM was required (Fig. 1h; Supplementary Fig. 3), as IgM monomers failed to clump the parasites (Fig. 1i, Supplementary Fig. 3). Of relevance, IgM-induced parasite clumping in eight diverse *Leishmania* species (Fig. 1j) strongly suggesting that parasite clumping upon contact with blood is a general response in *Leishmania*.

To directly establish the contribution of the IgM-induced clump to genetic exchange in vitro, we used 3 genetically marked *L. major* parasite strains with independent selectable markers (WR-SSU-HYG [parental line 1], FV1-FKP40-BSD [parental line 2], and FV1-FTL-SAT [parental line 3]) in various crosses. Stationary phase promastigotes from two parental lines were mixed at a 1:1 ratio and cultured for 3 days, with or without IgM, in the absence of selection drugs, and then grown in the presence of selective drugs for up to 30 days (Fig. 1k, l). Hybrids were observed for all parental combinations, but only when the culture was supplemented with IgM (Fig. 1k, l).

Collectively, our data demonstrate that IgM is critical for the generation of *L. major* hybrids in vitro through its induction of a structured and transient living parasite clump. Furthermore, without any other manipulation to the culture, in vitro hybrids from *L. tropica* crosses were only observed in the presence of IgM under our experimental conditions (Supplementary Fig. 4). Another breakthrough of this study was to have accomplished something sought after for many years, the formation of *L. major* hybrids in vitro.

Our studies addressed the hypothesis that formation of the IgM-dependent parasite clump promotes a closer contact between parasites, and this, in turn, increases the probability of *Leishmania* hybrid formation. Using confocal microscopy, we observed that IgM was abundantly distributed throughout the parasite clump in close association with *Leishmania* parasites, forming the distinct spherical structure observed in vitro (Fig. 2a). Transversal sections provided further evidence that *Leishmania* parasites lie in close proximity to each other within the parasite clump and revealed that IgM localized to distinct pockets between parasites (Fig. 2b) and coated their surface (Fig. 2c), potentially contributing to interactions holding the structure together. We then used scanning electron microscopy to establish the similarity in the organization of closely interacting parasites within the *L. major* (Fig. 2d, upper panel) and *L. tropica* (Fig. 2d, lower panel) clumps. Importantly, we located hybrid events inside a *L. tropica* parasite clump using two parental lines each harboring either a green or red nucleotide analog, only detected in the nucleus (Fig. 2e, left and middle panels). Double labeled nuclei (yellow), signifying a hybridization event, or a *Leishmania* hybrid, were only detected inside the parasite clump by confocal microscopy (Fig. 2e, right panel). Promastigotes with 3 nuclei (Fig. 2f, red arrows) indicative of fusion events were also observed inside the parasite clump by transmission electron microscopy. Our data provide direct evidence that IgM permeates the parasite clump and coats the parasites and establish that *Leishmania* fusion and hybrid formation take place within the parasite clump. Based on these findings and their relevance in parasite fusion and hybrid formation, we termed this structure the *Leishmania* mating clump (LMC).

**Fig 2.**
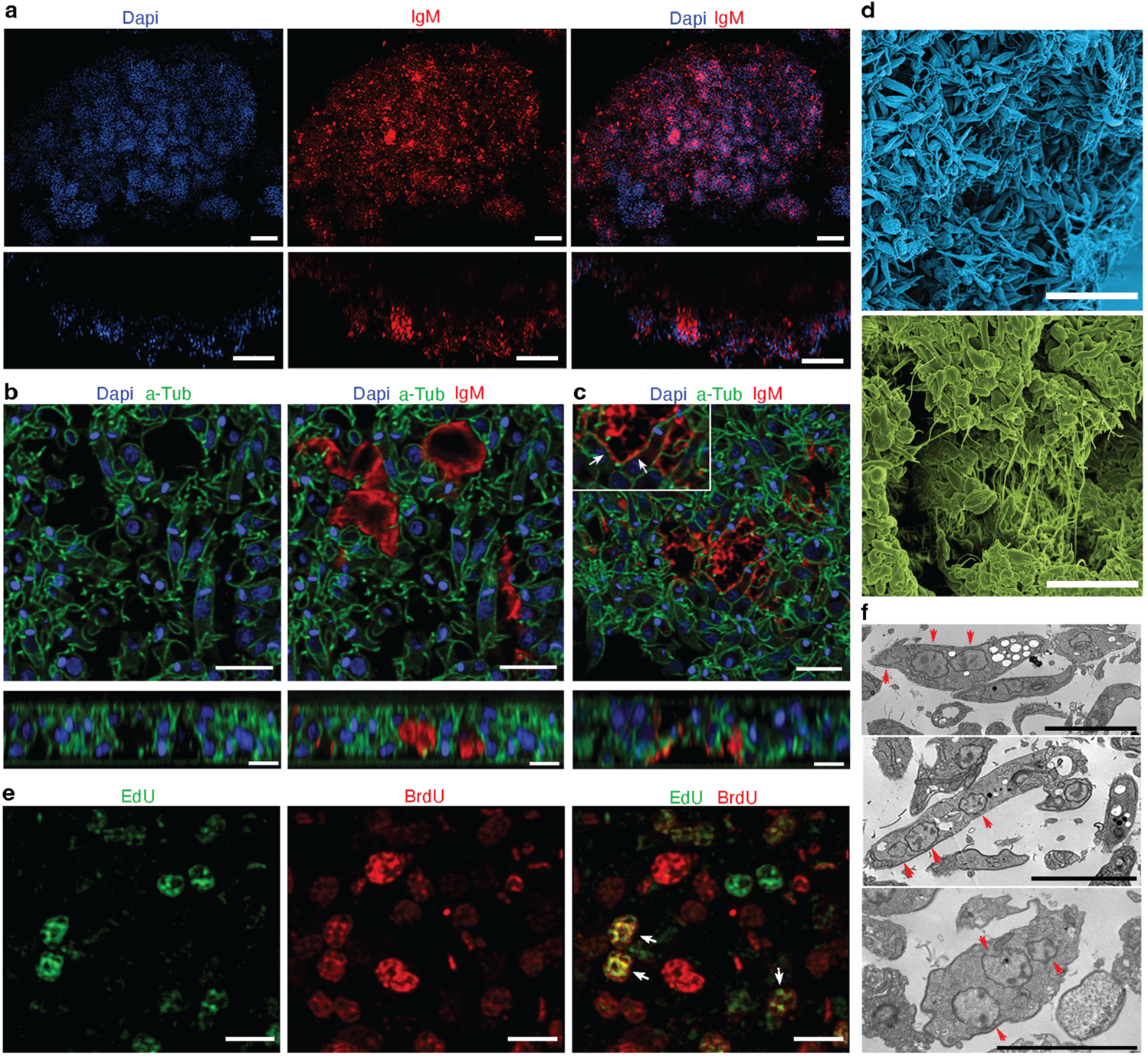
Characterization of the *Leishmania* mating clump. **(a)** Confocal immunofluorescence wide Z and tile images showing the association of IgM with the *L. major* mating clump after 24h of culture. Dapi (nuclei, kinetoplast) blue; IgM, red. Scale bar = 40 µm. Bottom rectangles for each panel represent XZ slices. Scale bar = 30 µm. **(b)** Confocal immunofluorescence of a 5 µm transversal section of the *L. major* mating clump. Dapi (nuclei; kinetoplast), blue; alpha-tubulin, green; IgM, red. IgM localizes between cells within the clump (right panel). Scale bars = 5 µm. Bottom rectangles for each panel represent XZ slices. Scale bars = 3 µm. **(c)** IgM colocalizes on the surface of parasites (arrows in inset). Scale bar = 5 µm. Bottom rectangles represent XZ slices. Scale bar = 3 µm. **(d)** SEM of a *L. major* (upper panel) and *L. tropica* (lower panel) mating clump showing the 3D organization of promastigote forms. Scale bar: 15 µm. **(e)** Confocal immunofluorescence of a 5 µm transversal section of the *L. tropica* mating clump. *L. tropica* promastigotes were grown to stationary phase in the presence of EdU or BrdU then extensively washed and used at 1:1 ratio to assemble the LMC with IgM. EdU, green; BrdU, red. White arrows point to yellow nuclei indicative of fusion and exchanged genetic material. Scale bars, 3 µm. (**f**) TEM of a *L. tropica* mating clump showing cells with 3 nuclei. Arrow, individual nucleus within a parasite cell body. Scale bar: 5 µm.

Eukaryotic class II fusogens such as HAP2, intrinsic molecules identified in *Chlamydomonas* and *Plasmodium*, are involved in membrane fusion between gametes. In contrast, IgM is an extrinsic molecule originating from the mammalian host that brings *Leishmania* parasites together to promote fusion, exposing a novel mechanism co-opted by a eukaryote to facilitate the early key steps of genetic exchange. We hypothesize that the LMC creates an environment, nutritional, physical, and biochemical that promotes parasite fusion and hybrid formation. It will be important to further investigate whether the interaction between IgM and the parasite is activating a signaling pathway, or whether the tight parasite scaffolding within the LMC is creating a physical or nutritional stress that facilitates genetic exchange. Nevertheless, it is important to emphasize that the addition of IgM to *Leishmania* culture media mimics natural exposure of *Leishmania* to IgM in sand fly blood meals. Every time a sand fly takes a blood meal, a *Leishmania* parasite that resides in the sand fly gut will interact with IgM present in the blood of all vertebrate animals^18–20^.

The ubiquity of IgM in vertebrate blood^18–21^ combined with the multiple and promiscuous (Supplementary Table 1) blood-feeding behavior of the sand fly^22^ led us to hypothesize that this antibody is a major facilitator for the initiation of genetic exchange in vivo, in the sand fly gut. Experiments were performed using conditions that mimicked the natural progression of sand fly infections, including incorporation of a second blood meal that is needed to prevent loss of naturally acquired parasites^22^. Having first established that IgM persists in the sand fly midgut for at least 24 hours after each blood meal (Fig. 3a), we performed an experiment where we allowed sand flies to feed on a cutaneous leishmaniasis lesion initiated by two *L. major* parental lines (Fig. 3b, BM1), to pick the parasites up as part of a natural blood meal ^23,24^. We then followed the natural sand fly feeding behavior by offering them a second uninfected blood meal six days after feeding on cutaneous lesions (Fig. 3b, BM2). The second blood meal was either devoid of IgM or supplemented with 500 µg/ml of IgM (Fig. 3b, BM2), a concentration at or below those reported in the blood of different animals (Supplementary Table 2). Additionally, a control group was maintained on a single infected blood meal (Fig. 3b, dotted arrow). Fourteen days after the initial infection, sand fly guts from each group were dissected and plated individually to screen for parasite hybrids (Fig. 3b). For the group initially infected with parental lines combination 1×2, IgM increased the proportion of hybrid formation by up to 12-fold in sand flies getting a second blood meal containing IgM compared to sand flies given a second blood meal devoid of IgM or sand flies given a single infectious blood meal (Fig. 3c, d, Supplementary Fig. 5a). In these two control groups, we observed a significantly lower proportion of hybrids as was previously reported^13,14^. The similar proportion of hybrids observed in flies given a second blood meal devoid of IgM or sand flies given a single infectious blood meal suggest that this may be a basal level of chance mating within the confined space of the sand fly midgut. It is worth noting that both control group scenarios would not occur in nature since an infected sand fly will always take a second blood meal 5-6 days after infection^1,22^. Strikingly, control groups lacking IgM did not generate any hybrids in parental lines combinations 1×3 and 2×3, further emphasizing the importance of IgM in *Leishmania* hybrid formation in vivo (Fig. 3c, d, Supplementary Fig. 5a). Of note, the parasite load, and percent metacyclics were similar in all experimental groups (Supplementary Fig. 5b-g). The pronounced effect of IgM on parasite hybrid formation when provided to sand flies six days after naturally acquired infections (BM2) suggests that proliferative stages are prone to genetic exchange. We argue that hybrids may form soon after a sand fly takes a blood meal but can only be detected later in the infection cycle after undergoing multiple cycles of division^1,23^. Collectively, the above data establish that IgM facilitates *Leishmania* hybrid formation inside the sand fly gut after naturally acquired infections, likely by bringing parasites closer together within a LMC that promotes fusion and hybrid generation. This is supported by the observation of structures similar to in vitro LMCs within the midgut (Supplementary Fig. 6; Supplementary Videos 7-12).

**Fig 3.**
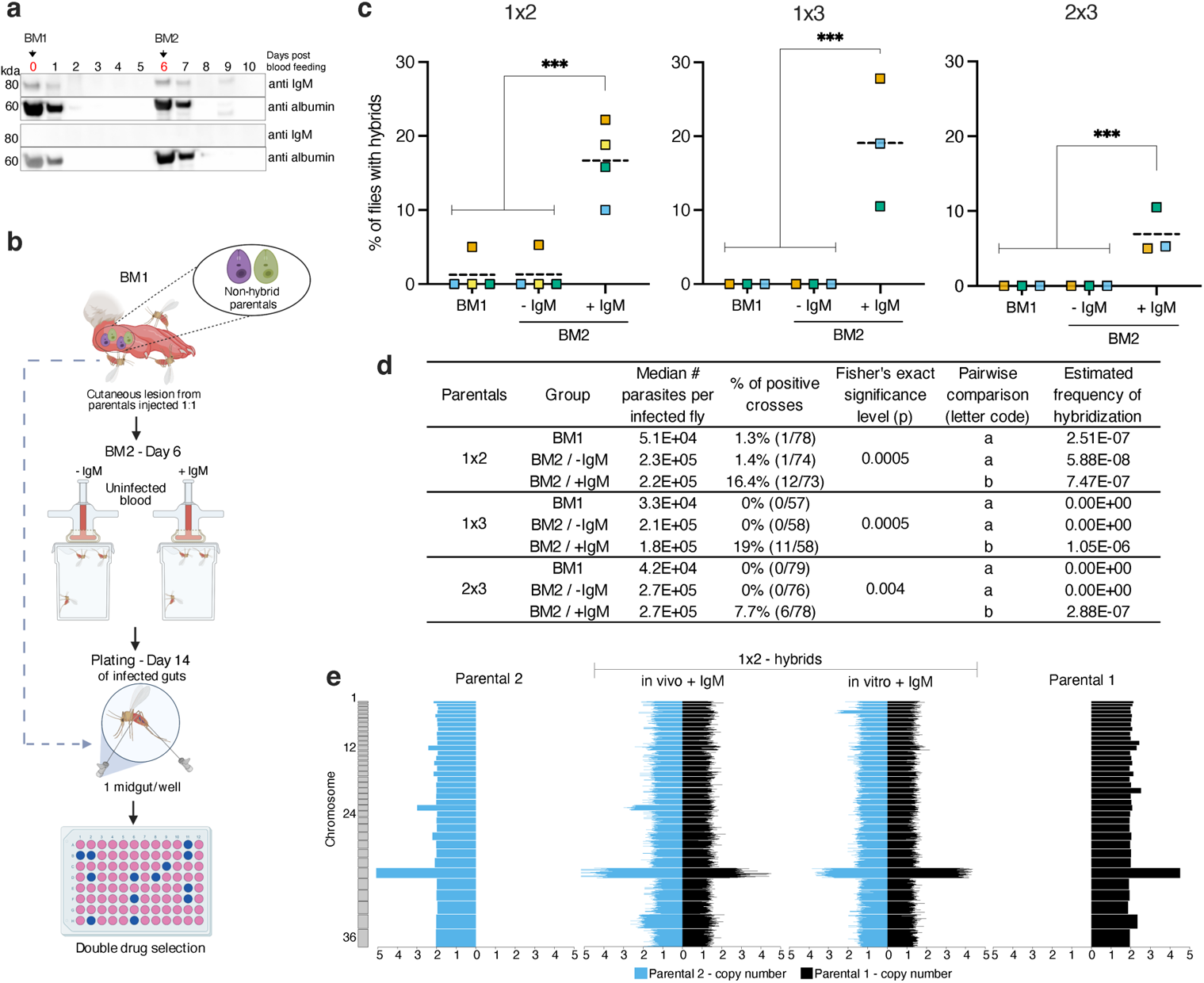
The impact of IgM on *Leishmania* hybrid formation in the sand fly vector. **(a)** Immunoblot detecting IgM in the sand fly midgut after multiple blood meals. Sand flies were provided an artificial uninfected blood meal with or without IgM. Arrows indicate the timing of the first (BM1, 0 days) and second (BM2, 6 days) blood meals. Pools of 15 midguts were used. 7.5 µg of total protein was loaded per lane. **(b)** Experimental design for the generation of in vivo crosses after sand flies acquire parasites by feeding on a footpad lesion containing two *L. major* parental lines. Bovine IgM was used. **(c)** Percent of sand flies harboring hybrids for 3 different combinations of parental lines (1×2; 1×3; 2×3) in single fed sand flies or in sand flies fed twice in the presence or absence of IgM. Colors indicate independent experiments. **(d)** Cumulative data from independent experiments for infection status and estimated frequency of hybridization per fly. 1×2 (n=4), 1×3 (n=3), 2×3 (n=3). Per each biological replicate ~ 20 infected flies per group. Fisher’s exact test, p < 0.05 is considered significant. Pairwise comparison indicate source of group significance: a, no difference; b, significant difference. **(e)** Whole-genome analysis of in vitro- and in vivo-generated hybrids for parental line combination 1×2. Biparental ancestry was confirmed across the whole genome for all crosses. Parental 1, WR-SSU-HYG; Parental 2, FVI-FKP40-BSD; Parental 3, FVI-FTL-SAT. *P < 0.05, **P ≤ 0.01, ***P ≤ 0.001.

Corroborating previous findings^10^, we observed hybrid formation only in sand flies and did not find them in *Leishmania* skin lesions or draining lymph nodes containing 2 different parasite lines (Supplementary Fig. 7). In the mammalian host, the *Leishmania* parasite is intracellular, residing in a parasitophorous vacuole inside macrophages^25,26^. It may be that a lack of opportunity for IgM to interact with amastigotes accounts for the absence of hybridization in the mammalian host. Whole-genome sequencing of the double drug resistant hybrids (Supplementary Fig. 8) showed that the crosses in sand flies and those generated in vitro were full genome hybrids (Fig. 3e; Supplementary Fig. 8-10). Moreover, aneuploidy commonly seen in cultured parasites^10,13,14^ were observed in this study at comparable levels (Supplementary Fig. 9, 10).

For use in positional cloning, generation of recombinant progeny by mating of hybrids to each other or to parental lines is required. Previous studies showed that the frequency of this progeny in *L. major* was considerably lower than it was for initial F1 hybrid formation, greatly hindering its utility^8^. To test whether insights from IgM-mediated mating could improve F1 hybrid recovery as well as confirm their fertility, we performed a backcross in the presence of IgM. This involved crossing a HYG/BSD double resistant (1×2) F1 hybrid with two clones of *L. major* parental line 2 bearing a SAT marker to allow identification of mating progeny. Sand flies were allowed to feed on a cutaneous lesion initiated by mixed infections of the (1×2) F1 hybrid and parental line 2’ or 2” of *L. major* (Fig. 4a, BM1), and then given a second uninfected blood meal six days later (BM2), either supplemented with 500 µg/ml of IgM or devoid of IgM (Fig. 4a, BM2). We consistently observed hybrid formation of this backcross, but only in sand flies fed with blood containing IgM (Fig. 4b, c; Supplementary Fig. 11a). Hybrid generation was not influenced by parasite density or percent metacyclics that were similar in all experimental groups (Supplementary Fig. 11b-g). A diagram outlining the approach to the generation of backcrosses is provided in Supplementary Fig. 12.

**Fig 4.**
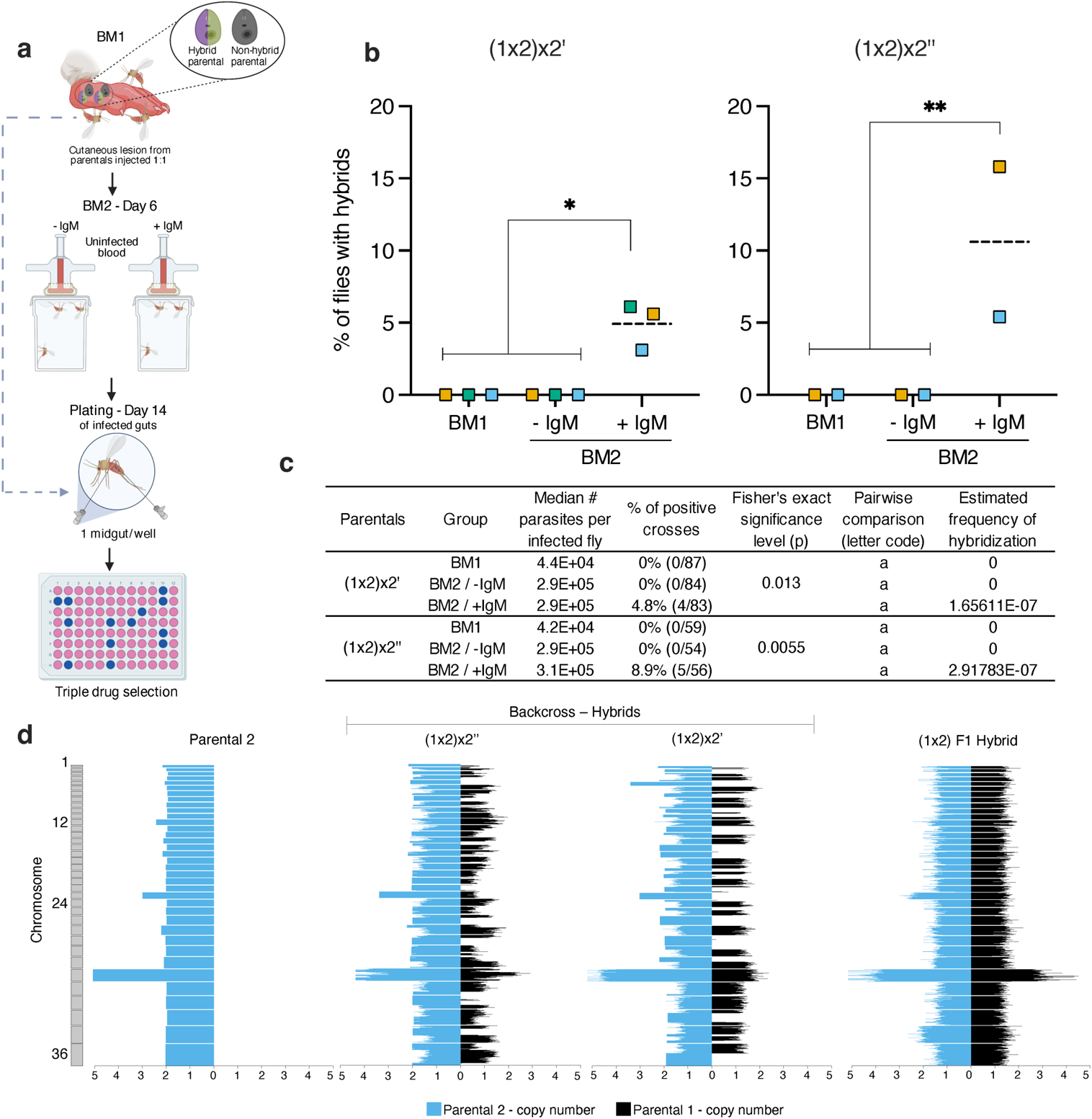
The impact of IgM on *Leishmania* genetic exchange in the sand fly vector. **(a)** Experimental design for the generation of in vivo backcrosses after sand flies acquire parasites by feeding on a footpad lesion containing one non-hybrid *L. major* parental line and one hybrid *L. major* parental line. A (1×2) F1 hybrid was injected at 1:1 proportion with the parental 2 harboring different resistance markers (parental 2’and parental 2”). Bovine IgM was used. **(b)** Percent of sand flies harboring hybrids in single fed sand flies or in sand flies fed twice in the presence or absence of IgM. Colors indicate independent experiments. **(c)** Cumulative data from independent experiments for infection status and estimated frequency of hybridization per fly. (1×2) × 2’ [n=3]; (1×2) × 2’’ (n=2). Per each biological replicate ~ 30 infected flies per group. Fisher’s exact test, p < 0.05 is considered significant. Pairwise comparison indicate source of group significance: a, no difference; b, significant difference. **(d)** Whole-genome analysis of in vivo-generated backcross hybrids for parental line combination 1×2. Where the biparental ancestry was consistent across each chromosome, recombination events have broken this up in the backcrosses, making parental copy number discontinuous always in the same direction through parental 2. Parental 1,WR-SSU-HYG; Parental 2, FVI-FKP40-BSD; Parental 2’, FVI-FKP40-SAT; Parental 2’’, FVI-SSU-SAT. *P < 0.05, **P ≤ 0.01.

Whole-genome sequencing of the triple drug resistant hybrids showed that the second-generation backcross progeny in sand flies bore the expected patterns of recombination between the F1, and 3^rd^ parent genomes also observed in previously reported backcrosses^8^. Importantly, no ‘backcross progeny’ were obtained with sand flies given a second blood meal devoid of IgM, providing conclusive evidence of its importance in *Leishmania* genetic exchange in a natural setting. Our data strongly suggest that IgM facilitates fertile hybrid formation and enhances their frequency in sand flies consequently promoting *Leishmania* diversity. Importantly, the reproducible and increased recovery of backcrosses opens the way for more in-depth genetic analysis of *Leishmania* parasites.

Historically, aggregation of *L. donovani* by IgM in culture was reported once as a standalone observation^27^. Our work provides direct experimental evidence of a critical role for IgM in LMC formation that promotes the first key steps of *Leishmania* genetic exchange in vitro and in vivo after a naturally acquired infection. Moreover, our data describe a new biological function for an antibody by implicating IgM in genetic exchange in a parasite. The exploitation of host IgM by *Leishmania* is of significant consequence. IgM is an evolutionarily conserved natural antibody that is always present in the blood of all vertebrates after birth^19–21^, independent of the immune status of the animal^28–33^. *Leishmania* evolved to exploit the pentameric properties of host IgM and the biological feeding pattern of sand flies to enhance the likelihood of mating. Sand flies ingest blood every 5-6 days to produce eggs^1^. We hypothesize that when a *Leishmania*-infected sand fly takes a subsequent blood meal, the periodic influx of IgM encounters parasites that have evolved to be at a stage that provides the density required to initiate the formation of the LMC in the confinement of the gut, a critical first step of genetic exchange that promotes fusion and hybrid generation. As such, in addition to metacyclic promastigotes, this implicates the two major proliferative stages, leptomonad and retroleptomonad promastigotes, as potential participants in the process of genetic exchange.

The fact that the *Leishmania* parasite co-opted a natural antibody from its mammalian host to advance its reproductive fitness uncovers a critical step in genetic exchange, providing an insight into the intimate evolutionary relationship between an insect vector, a parasite, and a mammalian host.

## Methods

### Ethics statement

All animal experimental procedures were reviewed and approved by the National Institute of Allergy and Infectious Diseases (NIAID) Animal Care and Use Committee under animal protocol LMVR4E. The NIAID DIR Animal Care and Use Program complies with the Guide for the Care and Use of Laboratory Animals and with the NIH Office of Animal Care and Use and Animal Research Advisory Committee guidelines. Detailed NIH Animal Research Guidelines can be accessed at https://oma1.od.nih.gov/manualchapters/intramural/3040-2/.

### Animals

Six to eight weeks old female BALB/c and Swiss Webster mice were obtained from Envigo and Charles River laboratories, respectively. Animals were housed under pathogen-free conditions at the NIAID Twinbrook animal facility, Rockville, MD.

### Parasites

*Leishmania* parasites used in this study: *Leishmania major* (WR 2885) isolated from a soldier deployed to Iraq^34^; *Leishmania major* (MHOM/IL/80/Friedlin); *Leishmania tropica* (MHOM/SU/74/K27 - ATCC 50129); *Leishmania braziliensis* (MHOM/BR/75/M2903 - ATCC 50135); *Leishmania infantum* (MCAN/BR/09/52) isolated from a dog spleen in Natal, Brazil^35^; *Leishmania donovani* (MHOM/SD/62/1S); *Leishmania tarentolae* (Tar II – ATCC 30143); *Leishmania mexicana* (MNYC/BZ/62/m379); *Leishmania amazonensis* (IFLA/BR/67/PH8). Parasites were grown at 27°C in Schneider’s Drosophila Insect Medium (Lonza BioWhittaker, 04-351Q) supplemented with 20% heat inactivated fetal bovine serum (FBS) (Gibco, 16140071) and 250 µM of adenosine (MilliporeSigma, A9251). Parasites were maintained by serial passage in BALB/c mice or Golden Syrian hamsters, depending on dermotropic or viscerotropic parasite species characteristics. Promastigotes were kept in culture for a maximum of 10 passages. *L. major* metacyclic promastigotes were purified by peanut agglutinin (PNA) (Vector Laboratories, L-1070) and used for in vitro experiments and infection of mice footpads as described^36^.

### *Leishmania* parental lines

The various *Leishmania* lines used in crossing experiments were constructed as detailed below. For parental line 1, *L. major* – WR-SSU-HYG/GFP, *L. major* (WR 2885) was used as background for generation of a hygromycin B (HYG) parental line that also harbors the EGFP gene. The EGFP gene was amplified by PCR using the mEGFP-C1 plasmid as a template for the forward primer 5-ACAGCGAGATCTATGGTGAGCAAGGGCGAGGA-3, and reverse primer 5-ACAGCGGGTACCCTTGTACAGCTCGTCCATGC-3. The forward and reverse primers add the BgIII or KpnI restriction sites to the amplicon, respectively. The PCR product was cloned into the Lexsy_hyg2 plasmid (Jena Bioscience, EGE-232). The linearized construct (2 µg) was used to transfect mid-log phase culture promastigotes on a 4D-NucleofectorTM X unit (Lonza) using pulse code FI-115 (T cell, Human, unistim. HE). Transfected parasites bearing rRNA locus-integrated EGFP were selected for growth in the presence of 100 µg/mL Hygromycin B (Goldbio, H270). Clones were selected by limiting dilution and assessed for chromosome integration. Parental line 2, *L. major* – FVI-FKP40-BSD^37^ and Parental line 3, *L. major* – FVI-FTL-SAT^38^ were generated as previously described. For generation of parental line 2’, *L. major* – FVI-FTL-SAT, we followed the procedures described for line 2, but used a heterozygous of the gene FKP40 with a construct harboring Nourseothricin (SAT). For parental line 2’’, *L. major* – FVI-SSU-SAT/RFP, *L. major* (MHOM/IL/80/Friedlin/FV1) was used as background for the generation of a SAT parental line that also harbors the mRuby gene. The mRuby gene was amplified by PCR using the mRuby-C1 plasmid as template for the forward primer 5-ACAGCGAGATCTATGAACAGCCTGATCAAAGA-3, and reverse primer 5-ACAGCGGCGGCCGCTCACAGATCCTCTTCAGAGATGAGTTTCTGCTCCCCTCCGCCCAGGCCGGC-3. The forward and reverse primers add the BgIII or NotI restriction sites to the amplicon, respectively. The PCR product was cloned into the Lexsy_sat2 plasmid (Jena Bioscience, EGE-234). The linearized construct (2 µg) was used to transfect mid-log phase culture promastigotes on a 4D-NucleofectorTM X unit (Lonza) using pulse code FI-115 (T cell, Human, unistim. HE). Transfected parasites bearing rRNA locus-integrated mRuby were selected for growth in the presence of 100 µg/mL Nourseothricin (Goldbio, N-500-100). Clones were selected by limiting dilution and assessed for chromosome integration. *L. tropica* (K27 – ATCC 50129) was used as background for the generation of parental line 4, *L. tropica* - K27-SSU-HYG/GFP and parental line 5, *L. tropica* - K27-SSU-SAT, respectively. Parental line 4 was made following the same steps as described for Parental line 1. Parental line 5 was made the same way as parental line 2’’ but without the addition of any fluorescent gene. mEGFP-C1 and mRuby-C1 were a gift from the Michael Davidson lab (Addgene plasmid # 54759, http://n2t.net/addgene:54759, RRID:Addgene_54759; Addgene plasmid # 54552, http://n2t.net/addgene:54552, RRID:Addgene_54552).

### In vitro *Leishmania* clumping activity

Various Leishmania species were cultured in complete Schneider’s medium in 25 cm^2^ flasks. 4×10^6^ /mL stationary phase promastigotes were dispensed into a well of a 24-well plate in the presence of 50 µg/mL of bovine IgM to test clumping activity. For *L. major*, sera from different animal sources were also tested for clumping activity in culture. Naïve beagle dog and germ-free mouse blood were provided by the Division of Veterinary Resources (DVR/NIH) under animal use protocol LMVR4E. Serum was obtained by centrifugation at 1000xg/10 min, inactivated at 56°C for 30 min, filtered sterilized through a 0.22 µm PES membrane (Millipore MILLEX GP, SGLP033RB) and stored at −80 in aliquots and used without thaw/freeze cycles. Sera acquired from commercial vendors are: Baboon (MilliporeSigma, S6884); Bovine - adult (MilliporeSigma, B9433); Bovine - fetal (Gibco, 16140071. R&D, S12495, S11595); Chicken (MilliporeSigma, C5405); Dog (BioChemed Services, TD092321NIH); Donkey (MilliporeSigma, D9663); Goat (MilliporeSigma, G9026); Guinea Pig (MilliporeSigma, G9774); Horse (MilliporeSigma, H1270); Human (MilliporeSigma, H4522, H6914, P2918, S7023, S1-M); Mouse (MilliporeSigma, S25-M, S3509, S7273); Porcine (MilliporeSigma, P9783); Rabbit (MilliporeSigma, R4505); Rat (MilliporeSigma, S7648); Sheep (MilliporeSigma, S2263). Sera were heat inactivated at 56°C for 30 min and filtered sterilized before usage at 20% final concentration.

### In vitro *Leishmania* mating clump assembly for recovery of hybrids

Parental lines of *L. major* were cultured in complete Schneider’s medium in 25 cm^2^ flasks. 3×10^6^ stationary phase promastigotes from each parental were mixed in a well of a 24-well plate with 1.5 mL of fresh complete Schneider’s medium + 50 µg/mL of bovine IgM, empirically determined as the optimal concentration that supports the formation and dissociation of the mating clump in vitro. Mating wells were maintained at 27°C for 3 days and then diluted 1:10 in fresh media in a 25 cm^2^ flask containing selective drugs. Cultures were followed for up to 30 days to allow for double-drug resistant cell lines to grow. Selective drugs were combined according to the parental lines used per experiment. The concentration used for each selective drug was as follows: Hygromycin B - 100 µg/mL (Goldbio, H270); Blasticidin - 10 µg/mL (Goldbio, B-800-500); Nourseothricin - 100 µg/mL (Goldbio, N-500-100). Once resistant cells grew, the polyclonal culture was cloned by plating 0.5 cell/well in a 96-well plate under double-drug pressure. After clone selection, genotyping was carried out using primers as displayed in Supplementary Table 3. Clone cultures were also a source of genomic DNA for deep sequencing. An illustration of the experimental design for in vitro *Leishmania* genetic exchange is provided in Fig. 1g. Frequencies of hybridization were estimated by: 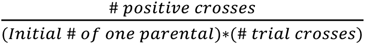

### Mice infection

Metacyclic promastigotes were harvested from stationary phase cultures and purified using peanut agglutinin (Vector Laboratories, L-1070) as previously described^36^. Purified *L. major* metacyclic promastigotes (1×10^5^) of each parental line were mixed at a 1:1 ratio and injected into a BALB/c mouse footpad. Animals that developed a swollen non-ulcerated (<5 mm thickness) footpad lesion at weeks 4 to 6 post-injection were used to feed sand flies. After sand fly feeding on infected animals, amastigotes were harvested from the infected footpads and skin draining lymph nodes as previously described^39^.

### Sand fly infection

Four to six days old female *Lutzomyia longipalpis* (Jacobina strain) sand flies were infected by feeding on mice footpad lesions of BALB/c mice. For first (F1) and second (backcross) generation crosses, culture purified metacyclic promastigotes of two parentals were injected in the mouse footpad at a 1:1 proportion. In F1 crosses experiments, mice were injected with parentals 1×2, 1×3 and 2×3. For backcross experiments mice were injected with a (1×2 F1 hybrid) × 2 harboring a different resistance marker. Illustration of the experimental design for natural feeding on mice is provided in Fig. 3b & 4a. Briefly, mice were anesthetized by intraperitoneal injection of a mixture of ketamine (100mg/kg) and xylazine (10mg/kg). The infected footpad was then inserted in a cardboard pint containing the sand flies. Sand flies were allowed to feed for 2 hours in the dark. After feeding, blood-fed female sand flies were sorted out and maintained at 26°C with 75% humidity and 30% sucrose solution *ad libitum*. *Lutzomyia longipalpis* is not the natural vector of *L. major* but it is a permissive sand fly species that has been shown to support the complete development of several *Leishmania* species from natural parasite acquisition to successful transmission by bite^40–42^. Importantly, midgut development related to parasites numbers and stages of differentiation in *Lu. longipalpis*/*L. major* infections was similar to those reported for *Phlebotomus papatasi / L. major* and *Lu. longipalpis / L. infantum* natural occurring combinations^43^.

### Preparation of the second blood meal provided to sand flies

For the 2^nd^ blood meal given six days post-infection to a proportion of the sand flies, defibrinated rabbit blood (Spring Valley Laboratories) was centrifuged at 1000xg for 10 min and the normal serum containing IgM was discarded. Red blood cells (RBCs) were washed three times with 10mL of PBS (Lonza BioWhittaker, BE17-516F) followed by centrifugation at 1000xg for 10 min. RBCs were washed an additional time with FBS (Gibco, 16140071). Blood was reconstituted by adding FBS at the same volume of the serum initially removed. Sand flies were allowed to feed for 2 hours on a glass feeder containing reconstituted blood supplemented or not with bovine IgM (MilliporeSigma, I8135) at 500 µg/mL. Unfed and blood-fed flies were sorted out after feeding and kept on 30% sucrose solution *ad libitum* for a total of 14 days after the initial infected blood meal. Females infected by feeding on mice lesions only were used to represent sand flies that have taken a single infected blood meal.

### Sand fly midgut counts

To assess maturity of the infection, *Leishmania*-infected sand fly midguts were individually transferred to a 1.7mL microtube (Denville Scientific, C2172) filled with 30μL of PBS. Midguts were then ground using a disposable pellet mixer and a cordless motor (Kimble, 7495400000). For counting, promastigotes were slowed down by a further 1:5 or 1:10 dilution in formalin/PBS at 0.005% final concentration. Then, 10 μL of the sample were loaded onto a Neubauer improved chamber (Incyto, DNC-NO1) and parasites were counted under a phase contrast Axiostar plus microscope (Zeiss) at 40X magnification. The number of parasites per midgut was back-calculated based on the total sample volume and subsequent dilution.

### In vivo recovery of hybrids

Only midguts of sand flies that developed a mature infection, recognized by an opaque anterior midgut distended by the presence of a mass of parasites^43^, were plated in search for hybrids. Infected midguts were dissected in PBS on microscope slides using tweezers and fine needles. Each midgut was transferred to a 1.7mL microtube (Denville Scientific, C2172) filled with 50 µL of complete Schneider’s medium and homogenized using a pestle with a cordless motor. One midgut was plated per well in a 96-well plate (Alkali Scientific, TP9096) at a final volume of 100 µL. An antibiotic cocktail (ABTs) that has no effect on parasite growth (Supplementary Fig. 13) and viability was included in complete media to control contamination with gut microbiota. ABTs: Penicillin-Streptomycin (Gibco, 100 U/mL); Gentamicin (Sigma, 50 µg/mL); Caspofungin (Sigma, 15 µg/mL); 5-fluorocytosine (Sigma, 30 µg/mL). For each independent experiment, a minimum of 20 infected midguts per group were collected. After plating, wells were allowed to stabilize and grow for 24h prior to the addition of 100 µL/well of complete Schneider’s media containing selective drugs (2x concentrated). Cultures were followed for up to 30 days for double or triple-drug resistant cell lines to grow. The concentration of selective drugs, cloning, and genotyping procedures were performed as described for in vitro recovery of hybrids. Frequencies of hybridization were estimated by: 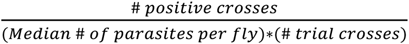.

### IgM identification from plasma by HPLC

Canine plasma (1.5-2 mL) was inactivated at 56°C for 30 min. Inactivated plasma was passed through a 0.22 µm PES membrane (Millipore MILLEX GP, SGLP033RB) and diluted to 7 mL in PBS containing 5 mM EDTA (buffer A). The inactivated, diluted plasma was then applied to a Sephacryl S200 (16/60, GE Healthcare) gel filtration column equilibrated with buffer A. Plasma components were eluted at 1 mL/min, and 2 mL fractions were collected and tested for *Leishmania* clumping activity. Active fractions were then combined, concentrated, and diluted several times with 10 mM sodium phosphate (pH 6.0) buffer (buffer B) before application to an SP Sepharose (16/10, GE Healthcare) ion exchange column equilibrated with buffer B. Proteins were eluted with a gradient of 0-1 M NaCl in buffer B. Activity was distributed throughout the major peak, particularly at the leading edge. These fractions were pooled and chromatographed on a Superdex 200 (10/300, GE Healthcare) gel filtration column equilibrated with buffer A, resulting in further purification of the major component that produced bands on SDS-PAGE gels of approximately 70 and 25 kDa consistent with the heavy and light chains of IgM. This sample was active in *Leishmania* clumping assays and was analyzed after tryptic digest by electrospray ionization mass spectrometry. As an additional means of purification, pooled fractions of IgM purified by gel filtration were diluted in 20 mM NaPO4 pH 7.5 (buffer C) containing 0.8 M ammonium sulfate and applied to a 1 mL HiTrap IgM purification HP column (GE healthcare) equilibrated with buffer C containing 0.8 M ammonium sulfate. The column was washed until the absorbance reached baseline levels. IgM was then eluted stepwise in buffer C and 0.5 mL fractions were collected. Activity in the *Leishmania* clumping assay closely aligned with the peak of absorbance.

### Commercial IgM preparation for in vitro and in vivo experiments

Commercially available IgM antibodies used in this study were obtained from MilliporeSigma and contained preservatives such as sodium azide. To avoid any potential toxicity to the parasites, removal of preservatives was undertaken. IgM from human (MilliporeSigma, I8260) and bovine (MilliporeSigma, I8135) sources were purified by passing them through a Sephacryl-S-200 gel filtration column (Cytiva) equilibrated with either buffer A minus EDTA or PBS. For IgM equilibrated with buffer A minus EDTA, further purification steps using a HiTrap IgM purification HP column are given above. Protein-containing fractions eluted with PBS were pooled and the volume reduced to the desired concentration by ultrafiltration.

### IgM stability in blood fed sand flies and its absence in fetal serum

To estimate how long IgM is maintained in the midgut after blood feeding, *Lu. longipalpis* were provided two uninfected blood meals on days 0 and 6, with or without 500 µg/mL of bovine IgM. Midguts were dissected daily on days 0 to 10 (*n=* 15) post initial blood meal and collected in 50µL of RIPA lysis buffer (Thermo-Fisher Scientific, 89901) supplemented with phenylmethylsulfonyl fluoride (PMSF) (MilliporeSigma, 93482) and Halt Protease Inhibitor Cocktail (Thermo-Fisher Scientific, 87786). Samples were gently macerated with a disposable pellet mixer and a cordless motor, then vortexed and centrifuged at 12,000 rpm for 10 min at 4°C. The supernatant was collected, and its protein content was quantified by BCA (Thermo-Fisher Scientific, 23227). A total of 7.5 µg of protein was loaded in a 15-well NuPAGE 4%–12% Bis-Tris protein precast gel (Thermo-Fisher Scientific, NP0335BOX), run under reducing conditions by the addition of the NuPAGE Sample Reducing Agent with DL-Dithiothreitol (DTT) (Thermo-Fisher Scientific, NP0009), and then denaturation at 95°C for 5 min. Proteins were transferred to a 0.45mm nitrocellulose iBlot 2 Transfer Stack membrane using the iBlot 2 Gel Transfer device (Thermo-Fisher Scientific, IB23001). To check for successful loading of antigens, No-Stain Protein Labeling Reagent (Invitrogen, A44717) was used following the membrane manufacture’s protocol. Next, membranes were blocked overnight at 4°C in 5% non-fat milk diluted in Tris-Buffered Saline, 0.1% TWEEN 20 (TBST) followed by 3 hours of incubation at RT with sheep anti-bovine IgM (Bio-Rad, AAI179) diluted at 1:1000. Membranes were washed three times with TBST followed by incubation for 30 min with rabbit anti-sheep IgG H+L secondary antibody conjugated to horse radish peroxidase (HRP) (Abcam, Ab6747) at a 1:5000 dilution. Membranes were washed three times with TBST and blots were developed by the addition of SuperSignal West Pico PLUS Chemiluminescence substrate (Thermo-Fisher Scientific, 34579). The signal was captured after 2 min using the Azure c600 chemiluminescence imager (Azure Biosystems). Membranes were stripped by adding 10 ml of Restore Western Blot Stripping Buffer (Thermo-Fisher Scientific, 21059) for 15 minutes at 37°C, followed by two washes with TBST. Blots were re-blocked overnight and incubated with recombinant rabbit anti-bovine serum albumin antibody (Abcam, ab192603) at 1:1000 dilution for 3 hours, followed by a 30 min incubation with HRP-conjugated goat anti-rabbit IgG H+L secondary antibody (Cell Signaling Technology, 7074S) at 1:1000 dilution. Blots were re-imaged as detailed above. To access the presence of IgM in fetal serum, NuPAGE 4%–12% Bis-Tris protein precast gels (Thermo-Fisher Scientific, NP0335BOX) were loaded with 1 µg of either Bovine IgM (Bio-Rad, AAI179), fetal bovine serum (Thermo-Fisher Scientific, 16140-071) or adult bovine serum (Millipore-Sigma, B9433) under reducing conditions followed by denaturation at 95°C for 5 min. The pattern of the proteins was visualized by Coomassie blue staining using the eStain™ L1 protein staining system. For immunoblotting, proteins were transferred onto nitrocellulose membranes, blocked, and probed with sheep anti-bovine IgM at 1:1000 followed by a 30 min incubation with HRP-conjugated anti-sheep IgG H+L secondary antibody (Abcam, Ab6747) diluted at 1:5000. Membranes were stripped and incubated with recombinant rabbit anti-bovine serum albumin antibody at 1:1000 followed by a 30 min incubation with HRP-conjugated goat anti-rabbit IgG H+L secondary antibody (Cell Signaling Technology, 7074S) at 1:1000 and imaged after addition of the chemiluminescence substrate.

### IgM depolymerization

To obtain monomeric IgM antibodies, 1 mg of bovine IgM (MilliporeSigma, I8135) was subjected to a digestion buffer exchange (50 mM Tris, 150 mM NaCl, 10 mM CaCl_2_, 0.05% NaN_3_; pH 8.0) - final concentration of 1 mg/ml. Afterward, 1 ml of IgM was incubated in the presence of 6 mg 2-mercaptoethylamine at 37 °C for 30 minutes in the dark. The reduced IgM was further alkylated in the presence of 16 mg iodoacetamide, followed by purification using a Superdex 200 Increase (10/300 GL; GE Healthcare) equilibrated with Buffer A minus EDTA. After size exclusion, a clear separation between the pentameric (peak 1 – P1) and monomeric (peak 3 – P3) IgM fractions were observed via native gel electrophoresis (NativePAGE Novex Bis-Tris Gel system; Life Technologies, Carlsbad, CA, USA). All fractions corresponding to defined peaks were pooled together, adjusted to a suitable concentration and frozen at −20 °C until further use.

### Mass spectrometry

Protein identification was carried out with a standard bottom-up method. The sample was reduced with DTT, alkylated with iodoacetamide, then digested using Lys-C followed by trypsin. The digested sample was cleaned using C18 tips (Agilent, OMIX10) and subjected to the LC-MS (EASY nLC 1200 and Orbitrap Fusion Lumos mass spectrometer, Thermo Scientific). The peptides were separated using a PepMap 100 C18 reversed phase column, and a standard data-dependent acquisition was performed. The survey MS1 (Top Speed mode, 3 seconds) was done using the Orbitrap mass analyzer, and the ddMS2 was done using the Linear Ion Trap where the fragmentation was done with CID. Dynamic exclusion was set for the duration of 20 seconds. The data analysis was done using PEAKS Studio software (Bioinformatcs Solutions Inc.). The database used is the UniProt proteome UP000002254 (Dog, taxonomy id 9615). The number of identified peptides, and protein abundance in the sample were confirmed by the PEAKS scores.

### Video recording and stereomicroscopic imaging

Midgut videos were taken using an iPhone 10 camera connected to the Stemi 508 stereomicroscope (Zeiss) ocular by a microscope mount (iDu Optics, LabCam iPhone 10 microscope adapter with built-in 30mm 10X WF lens). Phase contrast images and videos were recorded using a K5 camera coupled to a Thunder Imager microscope (Leica). Raw files were opened using Fiji ImageJ software^44^ and exported at 25 to 90 frames per second.

### Immunofluorescence of in vitro *Leishmania* mating clumps

After 24 hours of formation in vitro, whole Leishmania mating clumps were washed twice with PBS before fixation with 4% Paraformaldehyde (PFA) overnight at 4°C. Fixation was followed by permeabilization with 0.1% PBS Triton for 10 minutes, then 0.1 M glycine in PBS solution for 10 minutes, and a final blocking step with 0.1% Triton/0.1% BSA/ PBS for an hour at room temperature. Mating clumps were then incubated at 4°C overnight with mouse anti-alpha-tubulin monoclonal antibody (MilliporeSigma, T6199) conjugated to Alexa 555 (1:1000), and sheep anti-bovine IgM (Bio-Rad, AAI179) conjugated to Alexa 647 (1:1000) diluted in blocking buffer. Anti-Bovine IgM antibody was conjugated to an Alexa 647 fluorophore using the READILINK™ 647/674 Antibody Labeling Kit (Bio-Rad, 1351006) following the manufacturer’s instructions. The next day, mating clumps were washed three times with blocking buffer, ten minutes each, and then washed two times with permeabilization buffer, both at room temperature. Clumps were counterstained with 2 µM of Hoechst 33342 (Thermo Fisher Scientific, 62249) in permeabilization buffer for 30 minutes at room temperature, followed by a quick rinse of 0.1% Triton/PBS. Clumps were placed in a u-slide angiogenesis (IBIDI, 81506) well in Prolong gold antifade medium (Thermo Fisher Scientific, P10144) for imaging. For the sections, mating clumps were fixed in 4% paraformaldehyde overnight at 4°C, processed on the Leica ASP 6025, and then embedded in histology grade paraffin. All samples were sectioned at 3 µm. Mating clump section pretreatment was performed on the Leica Bond RX. Sections were baked for 30 minutes at 60°C and then dewaxed for 30 seconds in Bond Dewax solution (Leica, AR9222) heated to 72°C. Sections were rehydrated with absolute ethanol washes followed by 1x ImmunoWash (StatLab, ACR-015), then subjected to a 20-minute treatment with Epitope Retrieval Solution 1 (Leica, AR9961) heated to 100°C. Slides were then washed with PBS and stained. Steps for immunofluorescence of sections were based on Zaqqout et al. 2020^45^. Briefly, slides were permeabilized with PBS/gelatin/Triton 0.25%, twice for 10 minutes each at room temperature, followed by blocking with 5% BSA/PBS for one hour at room temperature in a humid box. The primary antibodies listed above were incubated in a blocking solution and kept overnight in a humid box at 4°C. The next day, slides were washed three times with PBS and then with permeabilization solution for 10 minutes at room temperature, and counterstained with 2 µM of Hoechst 33342 in the same buffer for 30 minutes at room temperature followed by a quick rinse with 0.1% Triton/PBS. Another final rinsing step with 10mM CuSO_4_/ 50mM NH_4_Cl solution for 10 minutes and a final short rinse with water was performed. Slides were closed using Prolong gold antifade mounting media and a coverslip.

### Confocal Microscopy

Images were captured using a Leica TCS SP8 DMi8 inverted fluorescence confocal microscope (Leica Microsystems) equipped with a photomultiplier tube/hybrid detector and a Leica DFC345 Monochrome camera. Images were taken using sequential acquisition, variable z-steps, mosaic size, and integration. Images were viewed with the 40x, 63x, and 100x oil immersion objective (zoom factor of 2, 3, or 4), and data were collected using the Leica Application Suite X software platform. White light laser and specific emission and excitation range were applied depending on the fluorophore used. The following spectra for excitation/emission were used: 488/520 for EdU and BrdU; 555/580 for alpha-tubulin; 594/620 for BrdU and EdU; and 647/665 for IgM. DAPI was excited using a 405-nm diode laser. Images were taken using sequential acquisition and variable z-steps. Whole mating clumps were imaged in the Z-wide mode (8-bit) and tile mode to capture the entire clump. Individual fields were merged and stitched together to mount the composite of the whole clump. Image processing was performed using Imaris 9.2.1 (Bitplane, Oxford Instruments). Maximum intensity projection (MIP) is represented in xy pictures, and cross-sections of between 2-3 µm were obtained with the ortho-slicer feature from Imaris.

### BrdU and EdU staining

*Leishmania* promastigotes were incubated with 10 µM of EdU - 5-Ethynyl-2’-deoxyuridine (Abcam, ab146186) or BrDu - 5-bromo-2’-deoxyuridine (Abcam, ab142567) for 4 days to reach the stationary growth phase. Parasites were then washed twice with PBS and an equal proportion of each of the labeled promastigotes was used for mating clump assembly. Mating clumps were fixed and processed for sections as described above. Three-micrometer sections were washed with PBS, permeabilized with 0.2% Triton/PBS for 10 minutes, then incubated with 0.1M glycine for 10 minutes at room temperature. Samples were then blocked with 0.5% BSA/PBS for 10 minutes and then briefly washed with PBS before the Click reaction was performed. We used the Click-it EdU Cell Proliferation Kit to image incorporated Edu with Alexa Fluor 488 dye (Invitrogen, C10337). The reaction was prepared following the manufacturer’s instructions and incubated with the samples for 30 minutes at room temperature in a humid and dark chamber. After the click reaction, sections were washed three times with PBS and blocked with 0.5% BSA/PBS for one hour at room temperature. Primary mouse monoclonal anti-BrdU (MOBU-1) (Invitrogen, B35128) (1:100) was incubated in blocking buffer overnight at 4°C. The next day, sections were washed with blocking buffer three times, 10 minutes each, and incubated with goat anti-mouse Alexa 594 secondary antibody (Invitrogen, A-11005) at 1:1000 in 0.1% Triton/PBS for 2 hours at room temperature. Slides were washed three times with 0.1%Triton/PBS for 10 minutes each and then counterstained with 2 µM of Hoechst 33342 in 0.1% Triton/ PBS for 30 minutes at room temperature. A final rinse was done with 10mM CuSO_4_/ 50mM NH_4_Cl solution for 10 minutes and followed by a quick rinse with water before the slides were closed with Prolong gold antifade mounting media and a coverslip.

### Genome sequencing

Genomic DNA was extracted from cloned parasite culture pellets using the E.Z.N.A.® DNA/RNA Isolation Kit (Omega Biotek, R6731) and following the manufacturer’s recommendations for cultured cells. Sample concentration was measured by a DS-11+ spectrophotometer (DeNovix Inc.). A minimum of 3 µg of DNA was sent for sequencing (Novogene Co.). Genomic DNA was randomly fragmented by sonication, then DNA fragments were end polished, A-tailed, and ligated with full-length adapters for Illumina sequencing followed by further PCR amplification with P5 and indexed P7 oligos. The PCR products were purified with the AMPure XP system and libraries were checked for size distribution using an Agilent 2100 Bioanalyzer (Agilent Technologies) and quantified by real-time PCR (to meet the criteria of 3 nM). Qualified libraries were fed into Illumina sequencers after pooling to obtain effective concentrations and the expected data volume.

### Genomic analysis

For Supplementary Fig. 8, raw sequencing reads were assessed for quality using the FastQC software. Fastq files were imported and processed using Geneious prime software v2021.2.2 (https://www.geneious.com). Low quality reads were trimmed from fastq files (< 20) and contaminating adapter primer sequences were removed using BBDuk trimmer plugin. Trimmed reads were mapped to the reference genome TriTrypDB-52_LmajorFriedlin using bowtie 2 plugin. BAM files with aligned reads were analyzed with qualimap v2.2.1^46^. The median genomic coverage of all samples was 33.1X. Using bowtie 2, unmapped reads were aligned to an artificial chromosome with selective markers. The artificial chromosome was created by concatenating the resistance genes integrated into the parental lines, HYG (Hygromycin), BSD (Blasticidin) and SAT (Nourseothricin), buffered by a 1.2kb (150bp repeated 8 times) sequence from mouse (*Mus musculus*) chromosome 1. For Fig. 3e, 4d and Supplementary Fig. 9, raw sequencing reads were trimmed using Trimmomatic v0.39^47^ and mapped using BWA-MEM v07.17^48^ to the TriTrypDB-52_LmajoprFriedlin reference genome with the resistant gene sequences inserted as additional contigs. PCR duplicates were marked using Samblaster v0.1.2.5^49^ and germline variants were called jointly across all samples using the HaplotypeCaller tool from GATK v4.1.9.0 and using default settings. Resulting variants were then filtered down to only biallelic sites with no missing data across all samples. The genomic insertion locations for each resistance gene were validated by examining split reads from each sample, and identifying the location of reads with one pair member mapped to the construct and the other pair member mapped to the human genome. Any locations with less than 3 supporting split read pairs were excluded from consideration. To quantify the allelic copy number of each contributing parent in F1 offspring, resulting BAM files were first used as input to the PAINT software suite^50^ to generate chromosome-level somy levels (Supplementary Fig. 10). A custom perl script was then used to execute the following steps with the multisample VCF and chromosome somy estimates as input: 1) identify homozygous SNP differences between parents for each cross, ignoring sites at <20X depth in both parental strains; 2) for each SNP identified in step 1, calculate the frequency of each allele based on read counts supporting each allele (e.g., allele1 proportion = (# reads supporting allele 1 at position1)/(total sequencing depth at position 1); 3) for a given F1, multiply the proportion of each allele by the chromosome-specific somy to get the copy number estimate for each parent. For the backcross hybrids analysis, the same procedure outlined above was used, and recombination breakpoints were identified as regions where parental copy number along a chromosome shifted by >=0.8 (to allow for 20% variance in absolute copy number estimation) across >=10 kilobases. All resulting copy number breakpoint calls were visually confirmed (e.g., Fig. 4d). The script used to execute the steps above is available here: https://github.com/jlac/Leishmania_allelic_copy_number.

### Scanning Electron Microscopy (SEM)

Leishmania mating clumps were fixed in 2.5% Glutaraldehyde in 0.1M Sodium Cacodylate Buffer, pH 7.4 (Electron Microscopy Sciences, 15960). Post fixation, the clumps were rinsed in 0.1M sodium cacodylate buffer and fixed with 1% OsO4 reduced with potassium ferrocyanide in buffer. After a couple of washes in water, the clumps were dehydrated in 100% ethanol and dried using multiple exchanges of hexamethyldisilazane (HMDS). On the following day, clumps were mounted on SEM pins using carbon adhesive tabs. Gentle pressure was applied to split open clumps to reveal their content, sputter coated with 5nm of Iridium using an EMS 300TD Quorum sputter coater, and imaged on a SU 8000 SEM.

### Transmission Electron Microscopy (TEM)

*Leishmania* mating clumps were fixed as per SEM protocol. After postfixing the organoids with 1% reduced OsO_4_ and dehydrating with 100% ethanol, the clumps were infiltrated and embedded with 100% Spurr’s resin in a flat mold. The flat mold with clumps were excised and mounted on BEEM™ capsules for trimming. Ultrathin sections were collected using a Leica UC7 ultramicrotome and imaged on an 80kV Hitachi 7800 TEM using an AMT™ bottom mount camera.

### Statistical analysis

All data manipulation and statistical analysis was performed in R (v. 4.1.1) with custom code written in Rstudio (v. 1.4.1717). Functions from the following packages were used in our scripts: abind^51^, chron^52^, data.table^53^, e1071^54^, emmeans^55^, FedData^56^, gsubfn^57^, gt^58^, ijtiff^59^, janitor^60^, lubridate^61^, MASS^62^, plyr^63^, rcompanion^64^, readxl^65^, reshape^66^, R.utils^67^, rlist^68^, Rmisc^69^, rstatix^70^, stats^71^, stringr^72^, tidyverse^73^ (contains: dplyr, ggplot2, forcats, tibble, readr, stringr, tidyr, purr), utils^71^, WRS2^74^, xlsx^75^. To compare hybrid formation success between the different experimental conditions (BM1, BM2/-IgM, BM2/+IgM) with the different parental line combinations (1×2, 1×3 & 2×3) as detailed in figures 3d and 4c, we applied an expanded Fisher’s exact test that could cope with r x n matrices over a chi-square test due to the occurrence of low expected values (<5). Since our data matrices exceed the standard 2×2 format, we followed the Fisher’s exact test with a pairwise test with the “pairwsieNominalIndependence” function from the R package rcompanion^64^ for 2-dimensional matrices, in which at least one dimension exceeded 2 levels. The output of this test was converted into a letter code using the “cldList” function from the R package rcompanion^64^ for improved test output representation; equal letters indicate no statistically significant difference, while different letters indicate a statistically significant difference (<0.05).

## Acknowledgments

We would like to thank Yvonne Rangel Gonzalez, Roberto Salas Carrillo, and Brittany Mills from VMBS, NIAID for technical support; Serena Doe and Diane Akueson from VMBS, NIAID for sand fly insectary support, Dr. Aline de Souza for critical reading of the manuscript. The authors thank Glenn A. Nardone, Motoshi Suzuki, and Lisa R. Olano of the Research Technologies Branch, NIAID, for mass spectrometry analysis. This research was supported by the Intramural Research Program of the NIH, National Institute of Allergy and Infectious Diseases, and NIH grant R01-AI29646 to S.M.B.

## End notes

### Author contribution

T.D.S. developed the hypothesis. T.D.S., S.K., J.G.V, S.M.B., I.V.C.A., C.B.M., J.A. and F.O. contributed to experimental design. E.I. planned the experiments. T.D.S., E.I., J.D., A.B.F.B., J.A., M.D., V.N., M.S., P.C. and T.W. performed the experiments. T.D.S., J.M.C.R, J.S.P.D, J.L., A.B.F.B., J.G.V., S.K., S.M.B. and E.I. Analyzed the data. C.M. performed sand fly insectary work. J.G.V., S.K. and S.M.B. supervised the project. All authors wrote the manuscript.

### Data availability

All data is present in the main text or the supplementary materials. Codes generated on this work used for copy number bioinformatics analysis on https://github.com/jlac/Leishmania_allelic_copy_number. Whole-genome sequencing raw reads files are deposited on SRA database under BioProject PRJNA848835. https://dataview.ncbi.nlm.nih.gov/object/PRJNA848835?reviewer=g3jklmr2bpu6ehm01aagjr68jv.

## Main References

1. Serafim, T. D. et al. Leishmaniasis: the act of transmission. Trends Parasitol 37, 976–987, doi:10.1016/j.pt.2021.07.003 (2021).

2. Burza, S., Croft, S. L. & Boelaert, M. Leishmaniasis. Lancet 392, 951–970, doi:10.1016/S0140-6736(18)31204-2 (2018).

3. Alvar, J. et al. Leishmaniasis worldwide and global estimates of its incidence. PloS one 7, e35671, doi:10.1371/journal.pone.0035671 (2012).

4. Tibayrenc, M. & Ayala, F. J. Is Predominant Clonal Evolution a Common Evolutionary Adaptation to Parasitism in Pathogenic Parasitic Protozoa, Fungi, Bacteria, and Viruses? Adv Parasitol 97, 243–325, doi:10.1016/bs.apar.2016.08.007 (2017).

5. Tibayrenc, M. & Ayala, F. J. The clonal theory of parasitic protozoa: 12 years on. Trends Parasitol 18, 405–410, doi:10.1016/s1471-4922(02)02357-7 (2002).

6. Tibayrenc, M., Kjellberg, F. & Ayala, F. J. A clonal theory of parasitic protozoa: the population structures of Entamoeba, Giardia, Leishmania, Naegleria, Plasmodium, Trichomonas, and Trypanosoma and their medical and taxonomical consequences. Proc Natl Acad Sci U S A 87, 2414–2418, doi:10.1073/pnas.87.7.2414 (1990).

7. Rogers, M. B. et al. Genomic confirmation of hybridisation and recent inbreeding in a vector-isolated Leishmania population. PLoS Genet 10, e1004092, doi:10.1371/journal.pgen.1004092 (2014).

8. Inbar, E. et al. Whole genome sequencing of experimental hybrids supports meiosis-like sexual recombination in Leishmania. PLoS Genet 15, e1008042, doi:10.1371/journal.pgen.1008042 (2019).

9. Van den Broeck, F. et al. Ecological divergence and hybridization of Neotropical Leishmania parasites. Proc Natl Acad Sci U S A 117, 25159–25168, doi:10.1073/pnas.1920136117 (2020).

10. Akopyants, N. S. et al. Demonstration of genetic exchange during cyclical development of Leishmania in the sand fly vector. Science 324, 265–268, doi:10.1126/science.1169464 (2009).

11. Stelzer, C. P. & Lehtonen, J. Diapause and maintenance of facultative sexual reproductive strategies. Philos Trans R Soc Lond B Biol Sci 371, doi:10.1098/rstb.2015.0536 (2016).

12. Kwok, A. J., Mentzer, A. & Knight, J. C. Host genetics and infectious disease: new tools, insights and translational opportunities. Nat Rev Genet 22, 137–153, doi:10.1038/s41576-020-00297-6 (2021).

13. Inbar, E. et al. The mating competence of geographically diverse Leishmania major strains in their natural and unnatural sand fly vectors. PLoS Genet 9, e1003672, doi:10.1371/journal.pgen.1003672 (2013).

14. Romano, A. et al. Cross-species genetic exchange between visceral and cutaneous strains of Leishmania in the sand fly vector. Proc Natl Acad Sci U S A 111, 16808–16813, doi:10.1073/pnas.1415109111 (2014).

15. Louradour, I., Ferreira, T. R., Ghosh, K., Shaik, J. & Sacks, D. In Vitro Generation of Leishmania Hybrids. Cell Rep 31, 107507, doi:10.1016/j.celrep.2020.03.071 (2020).

16. Louradour, I. et al. Stress conditions promote Leishmania hybridization in vitro marked by expression of the ancestral gamete fusogen HAP2 as revealed by single-cell RNA-seq. Elife 11, doi:10.7554/eLife.73488 (2022).

17. Walters, L. L. Leishmania differentiation in natural and unnatural sand fly hosts. J Eukaryot Microbiol 40, 196–206, doi:10.1111/j.1550-7408.1993.tb04904.x (1993).

18. Panda, S. & Ding, J. L. Natural antibodies bridge innate and adaptive immunity. J Immunol 194, 13–20, doi:10.4049/jimmunol.1400844 (2015).

19. Mashoof, S. & Criscitiello, M. F. Fish Immunoglobulins. Biology (Basel) 5, doi:10.3390/biology5040045 (2016).

20. Keyt, B. A., Baliga, R., Sinclair, A. M., Carroll, S. F. & Peterson, M. S. Structure, Function, and Therapeutic Use of IgM Antibodies. Antibodies (Basel) 9, doi:10.3390/antib9040053 (2020).

21. Boehm, T., Iwanami, N. & Hess, I. Evolution of the immune system in the lower vertebrates. Annu Rev Genomics Hum Genet 13, 127–149, doi:10.1146/annurev-genom-090711-163747 (2012).

22. Serafim, T. D. et al. Sequential blood meals promote Leishmania replication and reverse metacyclogenesis augmenting vector infectivity. Nat Microbiol 3, 548–555, doi:10.1038/s41564-018-0125-7 (2018).

23. Burkett-Cadena, N. D. in Medical and Veterinary Entomology (Third Edition) (eds Gary R. Mullen & Lance A. Durden) 17–22 (Academic Press, 2019).

24. Volfova, V. & Volf, P. The salivary hyaluronidase and apyrase of the sand fly Sergentomyia schwetzi (Diptera, Psychodidae). Insect Biochem Mol Biol 102, 67–74, doi:10.1016/j.ibmb.2018.09.010 (2018).

25. Chaves, M. M. et al. The role of dermis resident macrophages and their interaction with neutrophils in the early establishment of Leishmania major infection transmitted by sand fly bite. PLoS Pathog 16, e1008674, doi:10.1371/journal.ppat.1008674 (2020).

26. Lee, S. H. et al. Mannose receptor high, M2 dermal macrophages mediate nonhealing Leishmania major infection in a Th1 immune environment. J Exp Med 215, 357–375, doi:10.1084/jem.20171389 (2018).

27. Navin, T. R., Krug, E. C. & Pearson, R. D. Effect of immunoglobulin M from normal human serum on Leishmania donovani promastigote agglutination, complement-mediated killing, and phagocytosis by human monocytes. Infect Immun 57, 1343–1346, doi:10.1128/iai.57.4.1343-1346.1989 (1989).

28. Maddur, M. S. et al. Natural Antibodies: from First-Line Defense Against Pathogens to Perpetual Immune Homeostasis. Clin Rev Allergy Immunol 58, 213–228, doi:10.1007/s12016-019-08746-9 (2020).

29. Khasbiullina, N. R. & Bovin, N. V. Hypotheses of the origin of natural antibodies: a glycobiologist’s opinion. Biochemistry (Mosc) 80, 820–835, doi:10.1134/S0006297915070032 (2015).

30. Bendelac, A., Bonneville, M. & Kearney, J. F. Autoreactivity by design: innate B and T lymphocytes. Nat Rev Immunol 1, 177–186, doi:10.1038/35105052 (2001).

31. Hayakawa, K. et al. Positive selection of natural autoreactive B cells. Science 285, 113–116, doi:10.1126/science.285.5424.113 (1999).

32. Haury, M. et al. The repertoire of serum IgM in normal mice is largely independent of external antigenic contact. Eur J Immunol 27, 1557–1563, doi:10.1002/eji.1830270635 (1997).

33. Hayakawa, K., Hardy, R. R. & Herzenberg, L. A. Peritoneal Ly-1 B cells: genetic control, autoantibody production, increased lambda light chain expression. Eur J Immunol 16, 450–456, doi:10.1002/eji.1830160423 (1986).

## Methods references

34. Oliveira, F. et al. A sand fly salivary protein vaccine shows efficacy against vector-transmitted cutaneous leishmaniasis in nonhuman primates. Sci Transl Med 7, 290ra290, doi:10.1126/scitranslmed.aaa3043 (2015).

35. Aslan, H. et al. A new model of progressive visceral leishmaniasis in hamsters by natural transmission via bites of vector sand flies. J Infect Dis 207, 1328–1338, doi:10.1093/infdis/jis932 (2013).

36. Sacks, D. L. & Perkins, P. V. Identification of an infective stage of Leishmania promastigotes. Science 223, 1417–1419 (1984).

37. Guo, H. et al. Genetic metabolic complementation establishes a requirement for GDP-fucose in Leishmania. J Biol Chem 292, 10696–10708, doi:10.1074/jbc.M117.778480 (2017).

38. Vickers, T. J., Murta, S. M., Mandell, M. A. & Beverley, S. M. The enzymes of the 10-formyl-tetrahydrofolate synthetic pathway are found exclusively in the cytosol of the trypanosomatid parasite Leishmania major. Mol Biochem Parasit 166, 142–152, doi:10.1016/j.molbiopara.2009.03.009 (2009).

39. Afonso, L. C. & Scott, P. Immune responses associated with susceptibility of C57BL/10 mice to Leishmania amazonensis. Infect Immun 61, 2952–2959, doi:10.1128/iai.61.7.2952-2959.1993 (1993).

40. Cecilio, P. et al. Exploring Lutzomyia longipalpis Sand Fly Vector Competence for Leishmania major Parasites. J Infect Dis 222, 1199–1203, doi:10.1093/infdis/jiaa203 (2020).

41. Rogers, M. B. et al. Genomic confirmation of hybridisation and recent inbreeding in a vector-isolated Leishmania population. PLoS Genet 10, e1004092, doi:10.1371/journal.pgen.1004092 (2014).

42. Dostalova, A. & Volf, P. Leishmania development in sand flies: parasite-vector interactions overview. Parasit Vectors 5, 276, doi:10.1186/1756-3305-5-276 (2012).

43. Serafim, T. D. et al. Sequential blood meals promote Leishmania replication and reverse metacyclogenesis augmenting vector infectivity. Nat Microbiol 3, 548–555, doi:10.1038/s41564-018-0125-7 (2018).

44. Schindelin, J. et al. Fiji: an open-source platform for biological-image analysis. Nat Methods 9, 676–682, doi:10.1038/nmeth.2019 (2012).

45. Zaqout, S., Becker, L. L. & Kaindl, A. M. Immunofluorescence Staining of Paraffin Sections Step by Step. Front Neuroanat 14, 582218, doi:10.3389/fnana.2020.582218 (2020).

46. Okonechnikov, K., Conesa, A. & Garcia-Alcalde, F. Qualimap 2: advanced multi-sample quality control for high-throughput sequencing data. Bioinformatics 32, 292–294, doi:10.1093/bioinformatics/btv566 (2016).

47. Bolger, A. M., Lohse, M. & Usadel, B. Trimmomatic: a flexible trimmer for Illumina sequence data. Bioinformatics 30, 2114–2120, doi:10.1093/bioinformatics/btu170 (2014).

48. Li, H. Aligning sequence reads, clone sequences and assembly contigs with BWA-MEM. *arXiv preprint* arXiv:1303.3997 (2013).

49. Faust, G. G. & Hall, I. M. SAMBLASTER: fast duplicate marking and structural variant read extraction. Bioinformatics 30, 2503–2505, doi:10.1093/bioinformatics/btu314 (2014).

50. Shaik, J. S., Dobson, D. E., Sacks, D. L. & Beverley, S. M. Leishmania Sexual Reproductive Strategies as Resolved through Computational Methods Designed for Aneuploid Genomes. Genes (Basel) 12, doi:10.3390/genes12020167 (2021).

51. Plate, T. & Heiberger, R. abind: Combine Multidimensional Arrays. R package version 1.4-5. https://CRAN.R-project.org/package=abind (2016).

52. James, D. & Hornik, K. chron: Chronological Objects which Can Handle Dates and Times. R package version 2. 3–56. https://cran.r-project.org/web/packages/chron/index.html (2020).

53. Dowle, M. & Srinivasan, A. data.table: Extension of ‘data.frame’. R package version 1.14.0. https://CRAN.R-project.org/package=data.table (2021).

54. Meyer, D., Dimitriadou, E., Hornik, K., Weingessel, A. & Leisch, F. e1071: Misc Functions of the Department of Statistics, Probability Theory Group (Formerly: E1071), TU Wien. R package version 1.7-8. https://CRAN.R-project.org/package=e1071 (2021).

55. Lenth, R. V. emmeans: Estimated Marginal Means, aka Least-Squares Means. R package version 1.6.2-1. https://CRAN.R-project.org/package=emmeans (2021).

56. Bocinsky, R. K. FedData: Functions to Automate Downloading Geospatial Data Available from Several Federated Data Sources. R package version 2.5.7. https://CRAN.R-project.org/package=FedData (2019).

57. Grothendieck, G. gsubfn: Utilities for Strings and Function Arguments. R package version 0.7. https://CRAN.R-project.org/package=gsubfn (2018).

58. Iannone, R., Cheng, J. & Schloerke, B. gt: Easily Create Presentation-Ready Display Tables. R package version 0.3.1. https://CRAN.R-project.org/package=gt (2021).

59. Nolan, R. & Padilla-Parra, S. ijtiff: An R package providing TIFF I/O for ImageJ users. Journal of Open Source Software. 3(23), 663, doi:10.21105/joss.00633 (2018).

60. Firke, S. janitor: Simple Tools for Examining and Cleaning Dirty Data. R package version 2.1.0. https://CRAN.R-project.org/package=janitor (2021).

61. Grolemund, G. & Wickham, H. Dates and Times Made Easy with lubridate. Journal of Statistical Software, 40(3), 1–25. https://www.jstatsoft.org/v40/i03/ (2011).

62. Venables, W. N. & Ripley, B. D. Modern Applied Statistics with S. Fourth Edition. Springer, New York. ISBN 0-387-95457-0 (2002).

63. Wickham, H. The Split-Apply-Combine Strategy for Data Analysis. Journal of Statistical Software, 40(1), 1–29. http://www.jstatsoft.org/v40/i01/. (2011).

64. Mangiafico, S. rcompanion: Functions to Support Extension Education Program Evaluation. R package version 2.4.1. https://CRAN.R-project.org/package=rcompanion (2021).

65. Wickham, H. & Bryan, B. readxl: Read Excel Files. R package version 1.3.1. https://CRAN.R-project.org/package=readxl (2019).

66. Wickham, H. Reshaping Data with the reshape Package. Journal of Statistical Software 21, 1–20, doi:10.18637/jss.v021.i12 (2007).

67. Bengtsson, H. R. utils: Various Programming Utilities. R package version 2.10.1. https://CRAN.R-project.org/package=R.utils (2020).

68. Ren, K. rlist: A Toolbox for Non-Tabular Data Manipulation. R package version 0.4.6.1. https://CRAN.R-project.org/package=rlist (2016).

69. Hope, R. M. Rmisc: Rmisc: Ryan Miscellaneous. R package version 1.5. https://CRAN.R-project.org/package=Rmisc (2013).

70. Kassambara, A. rstatix: Pipe-Friendly Framework for Basic Statistical Tests. R package version 0.7.0. https://CRAN.R-project.org/package=rstatix (2021).

71. Team, R. C. R: A language and environment for statistical computing. R Foundation for Statistical Computing, Vienna, Austria. https://www.R-project.org/ (2021).

72. Wickham, H. stringr: Simple, Consistent Wrappers for Common String Operations. R package version 1.4.0. https://CRAN.R-project.org/package=stringr (2019).

73. Wickham, H. et al. Welcome to the tidyverse. Journal of Open Source Software, 4(43), 1686, https://doi.org/10.21105/joss.01686 (2019).

74. Mair, P. & Wilcox, R. Robust statistical methods in R using the WRS2 package. Behavior Research Methods 52, 464–488, doi:10.3758/s13428-019-01246-w (2020).

75. Dragulescu, A. & Arendt, C. xlsx: Read, Write, Format Excel 2007 and Excel 97/2000/XP/2003 Files. R package version 0.6.5. https://CRAN.R-project.org/package=xlsx (2020).

