## Supplementary Material for "Genetic exchange in *Leishmania* is facilitated by IgM natural antibodies"

### Supplementary information

Supplementary Figs. 1 to 12.

Supplementary Videos 1 to 12.

Supplementary Table 1 to 3.

Supplementary sequence 1.

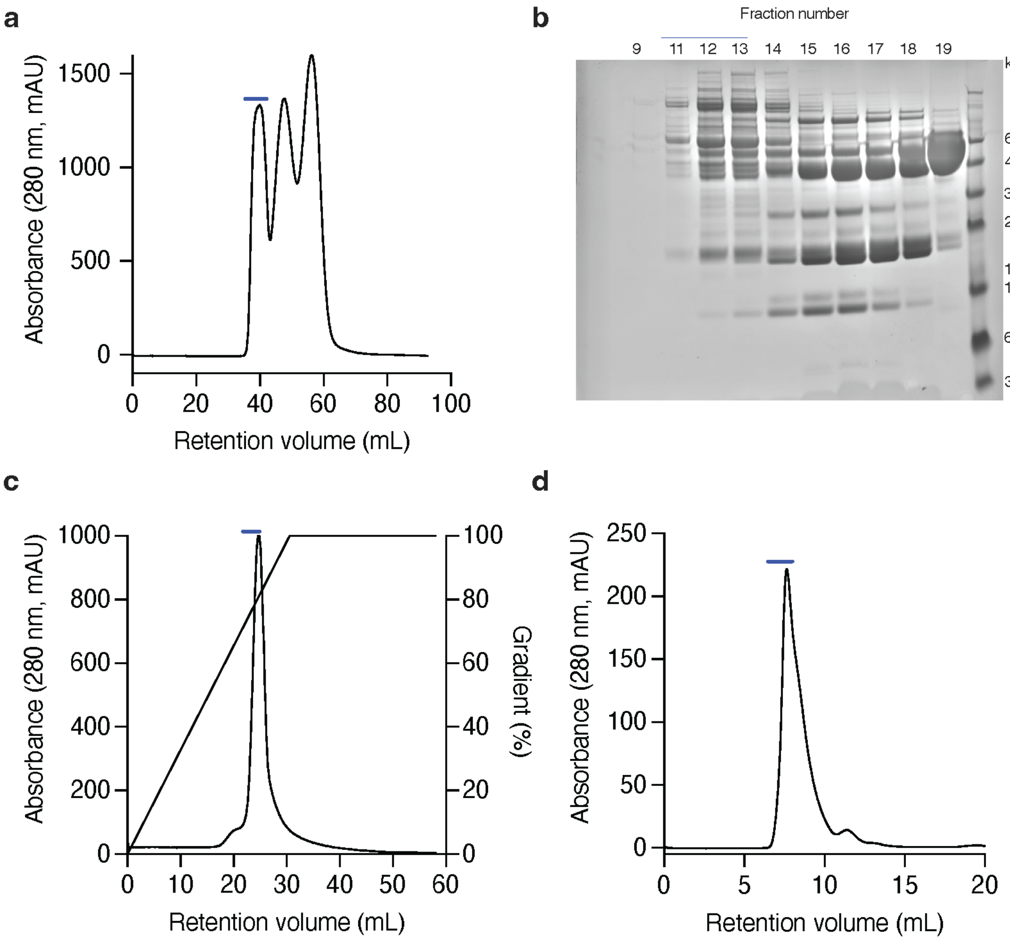


**e** Mass Spec. top hits

| Uniprot | Annotation | Area |
|---|---|---|
| J9NVC6 | Immunoglobulin heavy constant mu | 1.11E+10 |
| J9P9J6 | Immunoglobulin heavy constant mu | 6.05E+09 |
| J9JHH5 | Joining chain IgM, IgA | 3.00E+09 |
| H9GWR8 | Ig-like domain-containing | 2.89E+09 |
| F1PG16 | CD5-like | 2.22E+09 |
| J9NWX7 | Ig-like domain-containing | 6.19E+08 |
| P01874 | Ig mu chain C region | 3.12E+08 |
| F6UME0 | Alpha-2-macrogoblulin | 3.05E+08 |

**Supplementary Fig. 1. Identification of IgM in blood serum as the *Leishmania* clumping factor**. (**a**) Gel filtration chromatography of inactivated dog serum on Sephacryl S-200. Buffer: 20 mM sodium phosphate (pH 7.4), 150 mM NaCl, 5 mM EDTA. Bar represents HPLC fractions corresponding to *Leishmania* clumping activity. (**b**) SDS-PAGE of fractions obtained from the gel filtration chromatography shown in panel A. Bar represents fractions containing clumping activity that were then pooled for ion exchange chromatography. (**c**) Ion exchange chromatography of pooled fractions 11-13 from B. HPLC gradient, a sodium chloride gradient of 0 – 1 M in 20 mM sodium phosphate, pH 6.0. Bar represents fractions pooled for high resolution gel filtration chromatography. (**d**) High-resolution gel filtration chromatography on Superdex-200. Buffer: 20 mM sodium phosphate (pH 7.4), 150 mM NaCl. Bar represents fractions pooled for protein identification. (**e**) Mass spectral analysis of the most abundant proteins in pooled fractions from d.

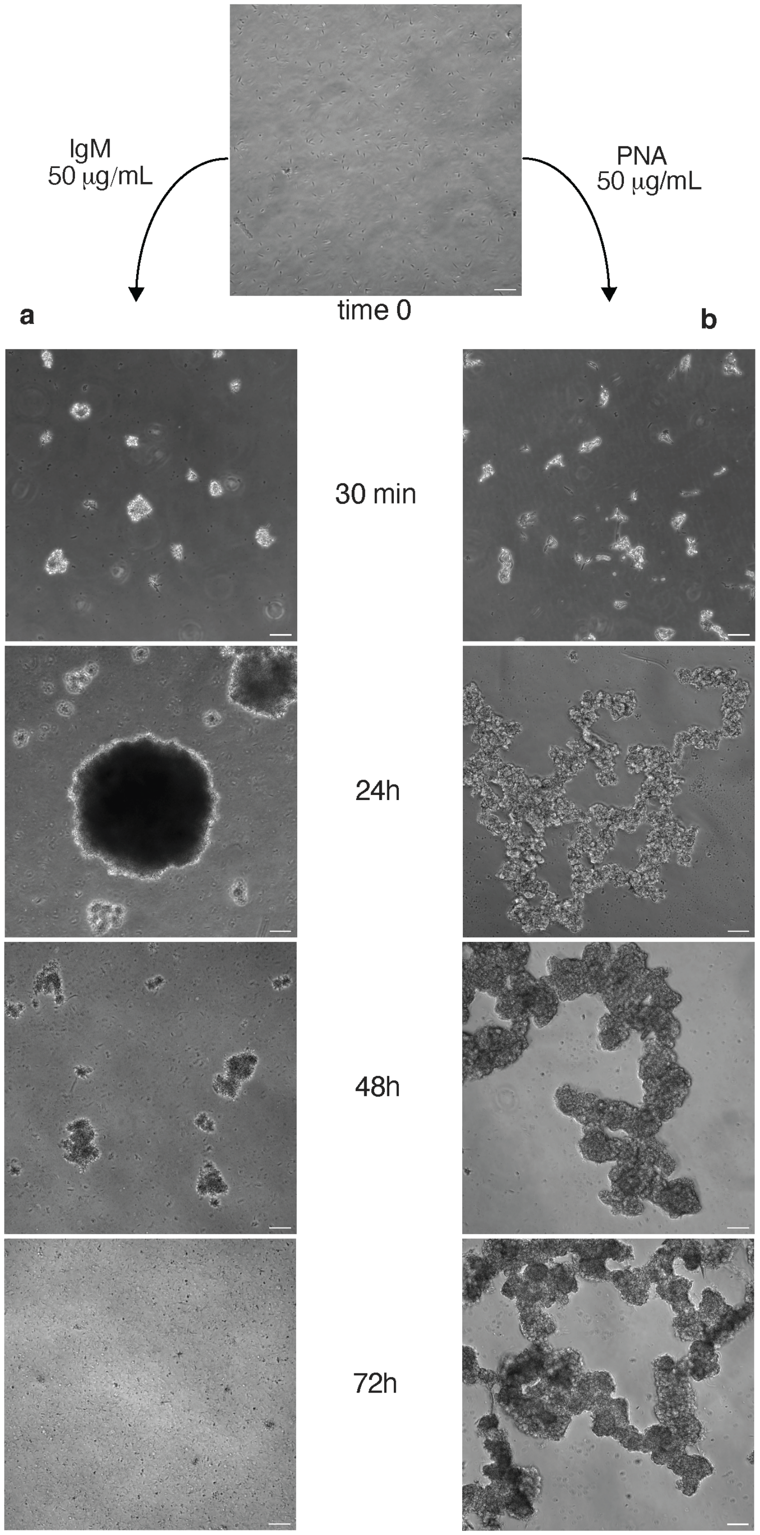


**Supplementary Fig. 2. IgM-induced clump formation differs from PNA agglutination**. *Leishmania major* promastigotes were grown to stationary phase and used under new media condition for clump pattern evaluation. **(a)** Parasites incubated with bovine IgM and imaged over the course of 72 hours. Cells rapidly start to clump, and individual clumps keep fusing for 24 hours. After 36 hours a gradual dissociation begins. By 72 hours, most of clumps are fully dissociated. Parasites in the clumps remain highly mobile throughout the process. **(b)** Parasites from the same original culture were incubated with peanut agglutinin (PNA) over the course of 72 hours. The agglutination pattern is distinct from the IgM-induced clumping and to the extent of the experimental timeline, it was irreversible. Agglutinated parasites were sluggish with low or no mobility. Scale bars = 50 μm.

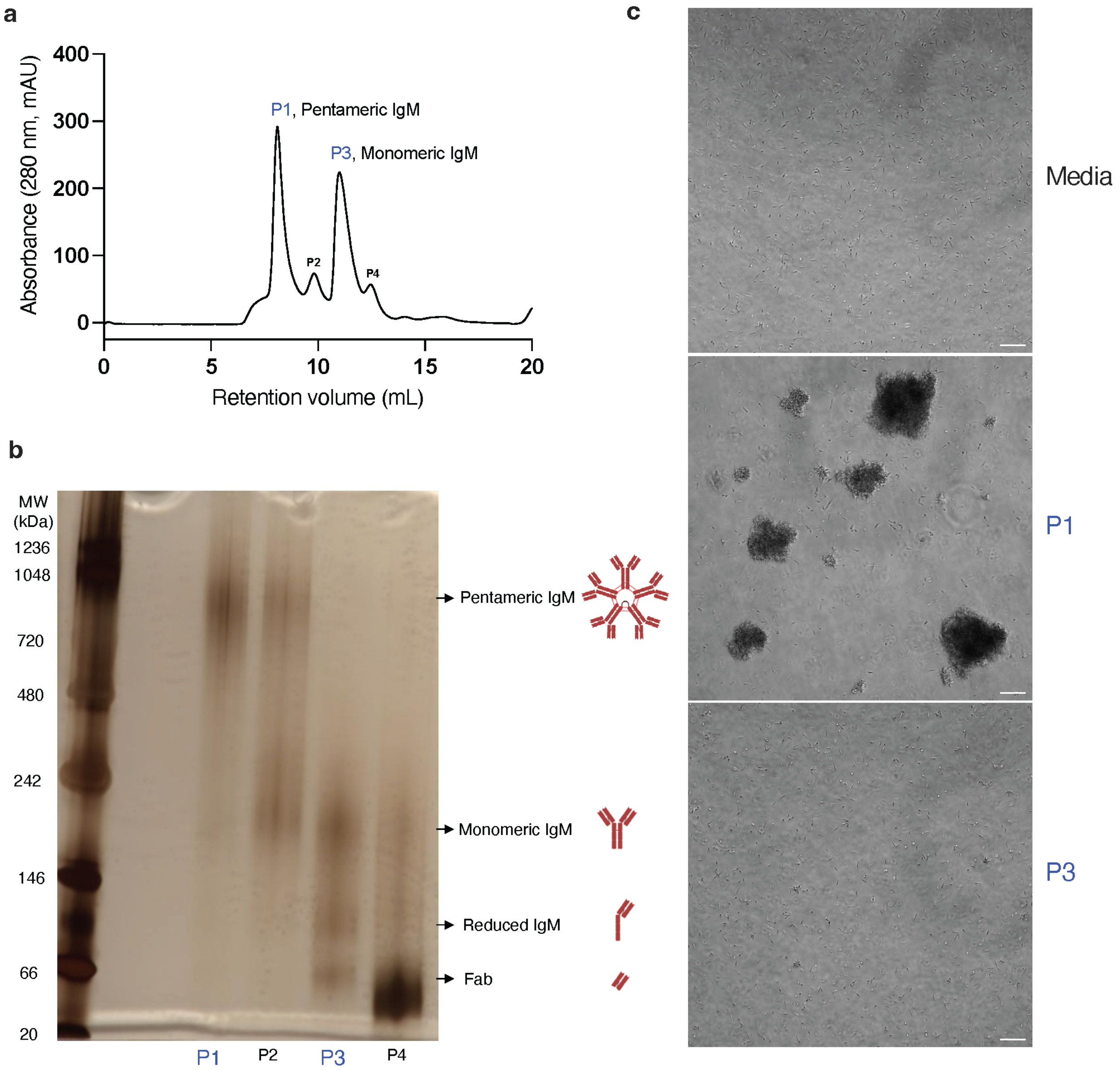


**Supplementary Fig. 3. IgM-Induced clumping depends on the multimeric nature of the antibodies.** (**a**) High-resolution gel filtration chromatography of IgM antibodies on a Superdex-200 Increase column after reduction and alkylation. Buffer: PBS. Four peaks. P1 to P4, are visible. **(b)** Native gel electrophoresis of the fractions corresponding to peaks P1 to P4 after staining with silver (silver staining kit – Thermo Scientific 24612). Bands with molecular weights suggestive of pentameric, monomeric, and reduced IgM antibodies, as well as the IgM Fab fraction are highlighted. **(c)** Phase contrast images of *L. major* promastigotes in culture media containing 20% fetal bovine serum alone (Media), or with 50 µg/ml of either control IgM (P1) or purified monomeric IgM after reduction/alkylation (P3). Images were taken 2 h after cells were seeded in fresh media. Scale bar = 50 µm.

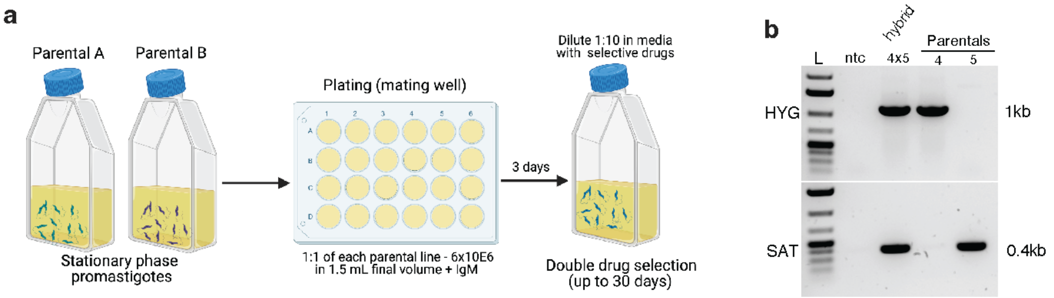


c

| Parentals | Initial # of each parental cells | # of trial crosses | Group | Clump formation | % of positive crosses | Estimated frequency of hybridization |
|---|---|---|---|---|---|---|
| 4x5 | 3.0E+06 | 12 | - | - | 0% (0/12) | 0.0E+00 |
| | | | IgM | + | 33.3% (4/12) | 1.1E-07 |

**Supplementary Fig. 4. IgM promotes *Leishmania tropica* hybrid formation in vitro. (a)** Workflow for in vitro crossing of *L. tropica* parental lines. Mating wells were established with 50 µg/mL IgM in complete Schneider's media. Double drug resistant hybrid parasites were cloned before genotyping. (**b**) *Leishmania* hybrids genotyping by PCR targeting parental selectable drug markers HYG (Hygromycin), and SAT (Nourseothricin). Parental 4, *L. tropica* - K27-SSU-HYG; Parental 5, *L. tropica* - K27-SSU-SAT; ntc, no template control; L, 1kb plus ladder (Invitrogen). (**c**) Summary of data from in vitro crossing of parental lines 4x5 (n=12).

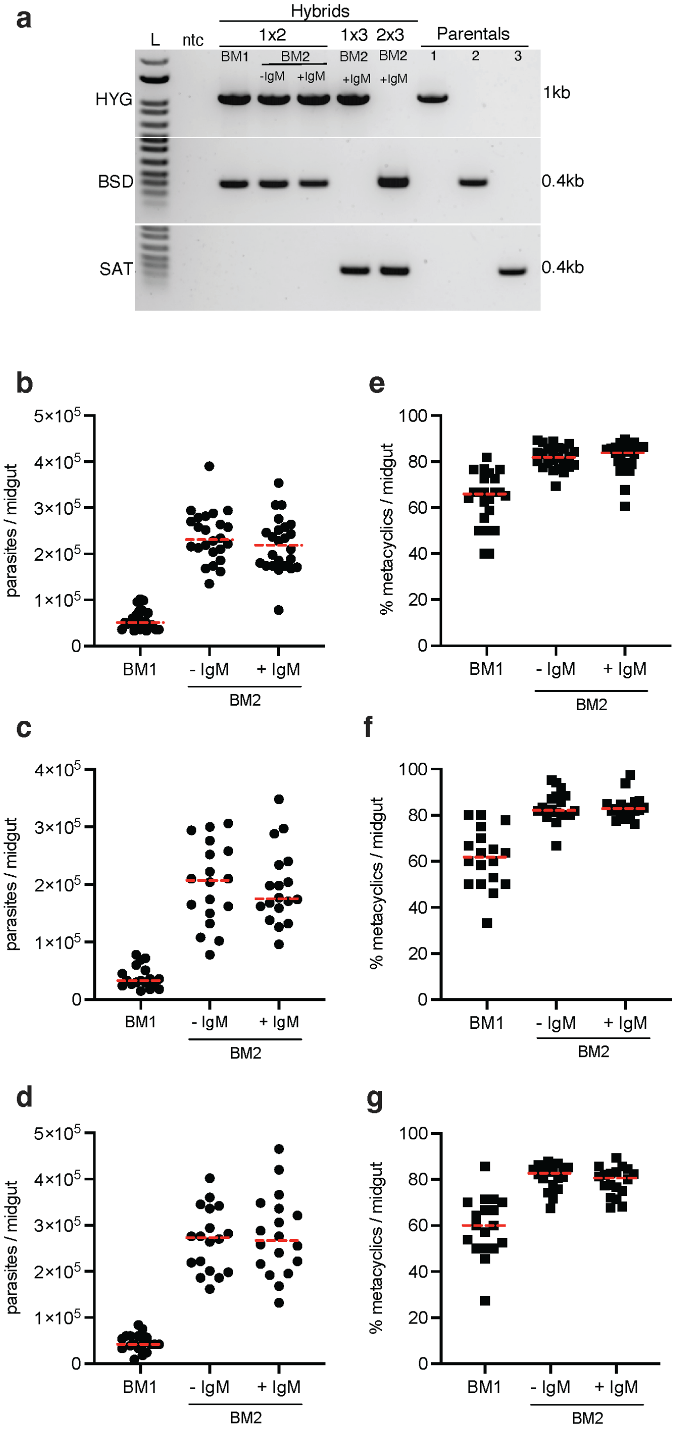


**Supplementary Fig. 5. Hybrid genotyping and infection status in sand flies after a naturally acquired *Leishmania* infection from mice lesions composed of two parental lines. (a)** *Leishmania major* hybrids formed in sand flies given one infectious blood meal (BM1) or provided

6 days later with an additional uninfected bloodmeal (BM2) in the presence (+IgM) or absence (-IgM) of IgM were genotyped by PCR targeting parental selectable drug markers HYG (Hygromycin), BSD (Blasticidin) and SAT (Nourseothricin). Parental 1, WR-SSU-HYG; Parental 2, FVI-FKP40-BSD; parental 3, FVI-FTL-SAT; ntc, no template control; L, 1kb plus ladder (Invitrogen). Double drug resistant hybrid lines were cloned before genotyping. A single hybrid representative is shown for positive events from each group. Parasite number (**b,c,d**) and percentage of metacyclic promastigotes (**e,f,g**) in *L. major*-infected *Lu. longipalpis*. At 6 days post-infection, a proportion of the sand flies were provided a second uninfected blood meal. Infection status of individual sand flies was assessed at 14 days after the first blood meal or 8 days after the second bloodmeal. Parental line combination 1x2 (b,e), 1x3 (c,f) and 2x3 (d,g).

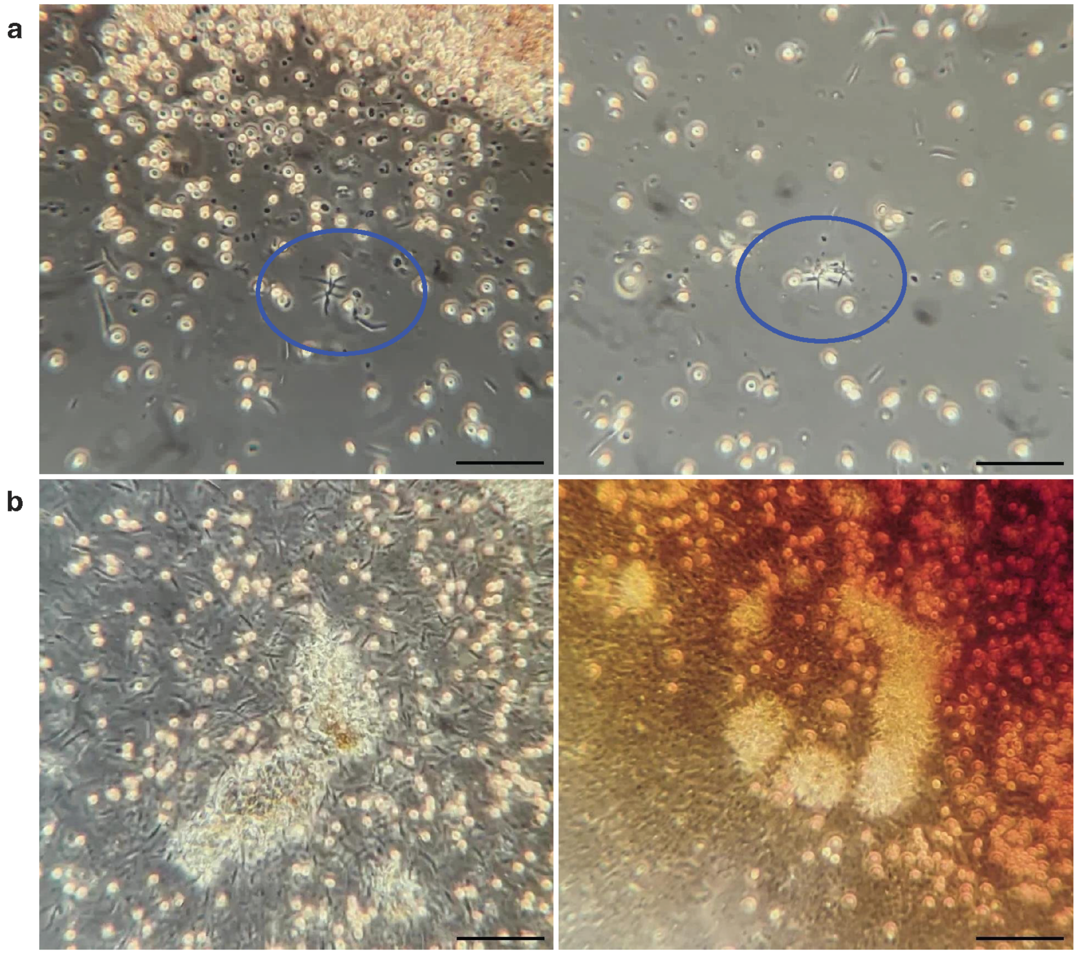

**Supplementary Fig. 6. *Leishmania* mating clump formation within the sand fly midgut.** Twelve days post-infection with *L. major*, *Lu. longipalpis* females were provided with a second blood meal containing IgM (500 µg/mL). **(a)** Initiation of *Leishmania* mating clump (LMC) formation 30 min after imbibing a second blood meal (blue circles). **(b)** Parasites fuse to form larger LMCs within 24 hours after imbibing a second blood meal. LMCs fusion in sand flies can also be observed in Supplementary Videos 11 and 12. Scale bars = 50 µm.

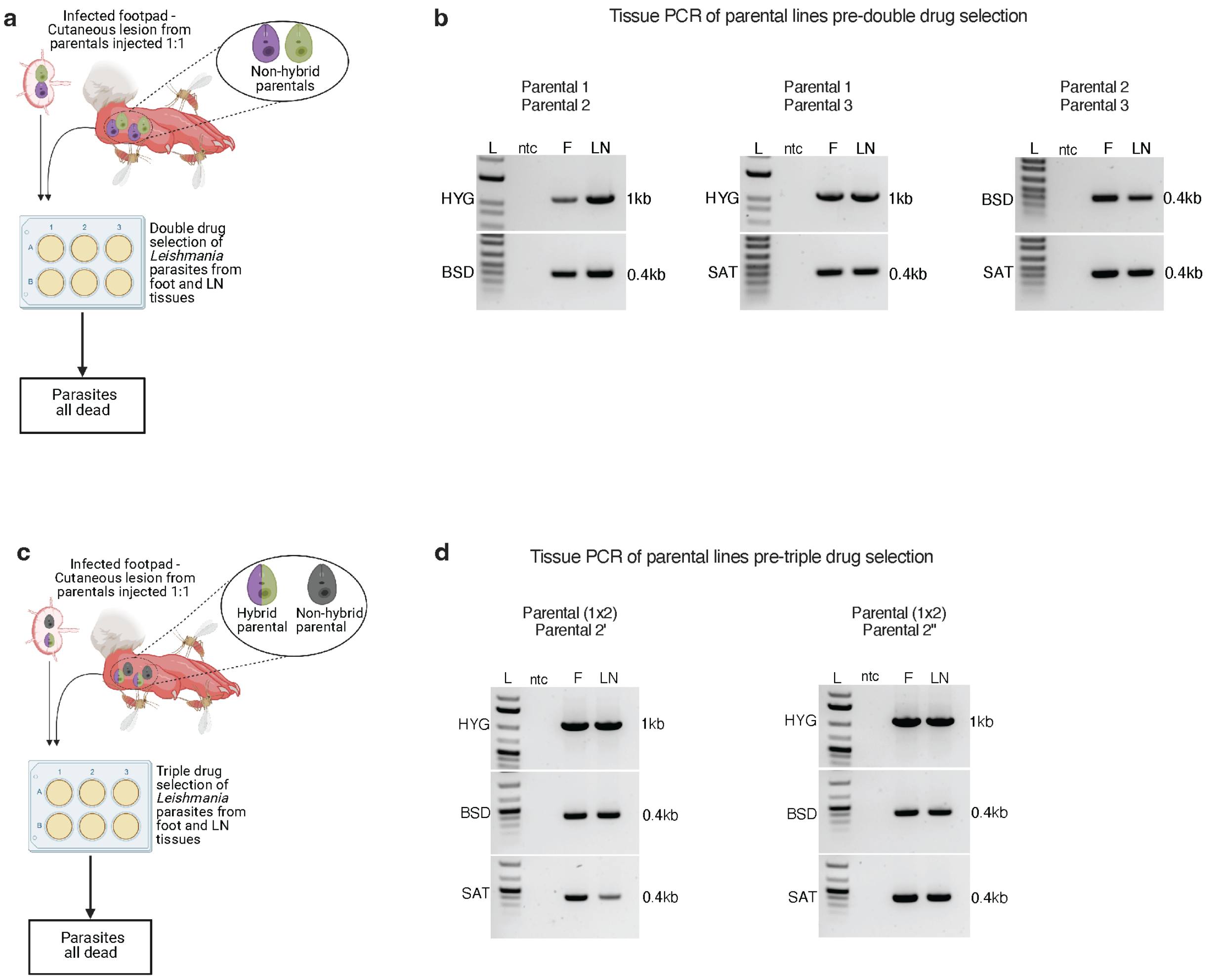


**Supplementary Fig. 7. Evaluation of parental lines and hybrids in infected mouse tissue. (a)** Experimental design. Mouse footpads were injected with a 1:1 mixture of two *L. major* parental line combinations 1 and 2, 1 and 3, or 2 and 3. After lesion development, 3 to 5 weeks post injection, infected footpads were exposed to sand flies for their first blood meal. Tissue from the infected footpad and draining lymph node (LN) were then disrupted and seeded in complete Schneider's media (6 mL) for 3 days to allow for amastigote differentiation into promastigotes. Half of the material was then cultured in double drug pressure for selection of hybrids. (**b**) The second half of the tissue from (**a**) was used for DNA extraction to confirm the presence of both parental lines by genotyping. (**c**) Experimental design. Mouse footpads ware injected with a 1:1

mixture of F1 hybrid of 1x2 and the parental 2 harboring a different resistance marker (SAT) at the same (2') or different (2'') chromosome locus. Sand fly feeding on mouse lesions and tissue processing were carried out as outlined above. Half of the tissue from the footpad and draining lymph node was then cultured in triple drug pressure for selection of hybrids. (**d**) The second half of the tissue from (**c**) was assessed for the presence of both parental lines by genotyping. Parental 1, WR-SSU-HYG; Parental 2, FV1-FKP40-BSD; Parental 3 = FV1-FTL-SAT; Parental 2', FV1-FKP40-SAT; Parental 2'', FV1-SSU-SAT.

Supplementary Fig.8

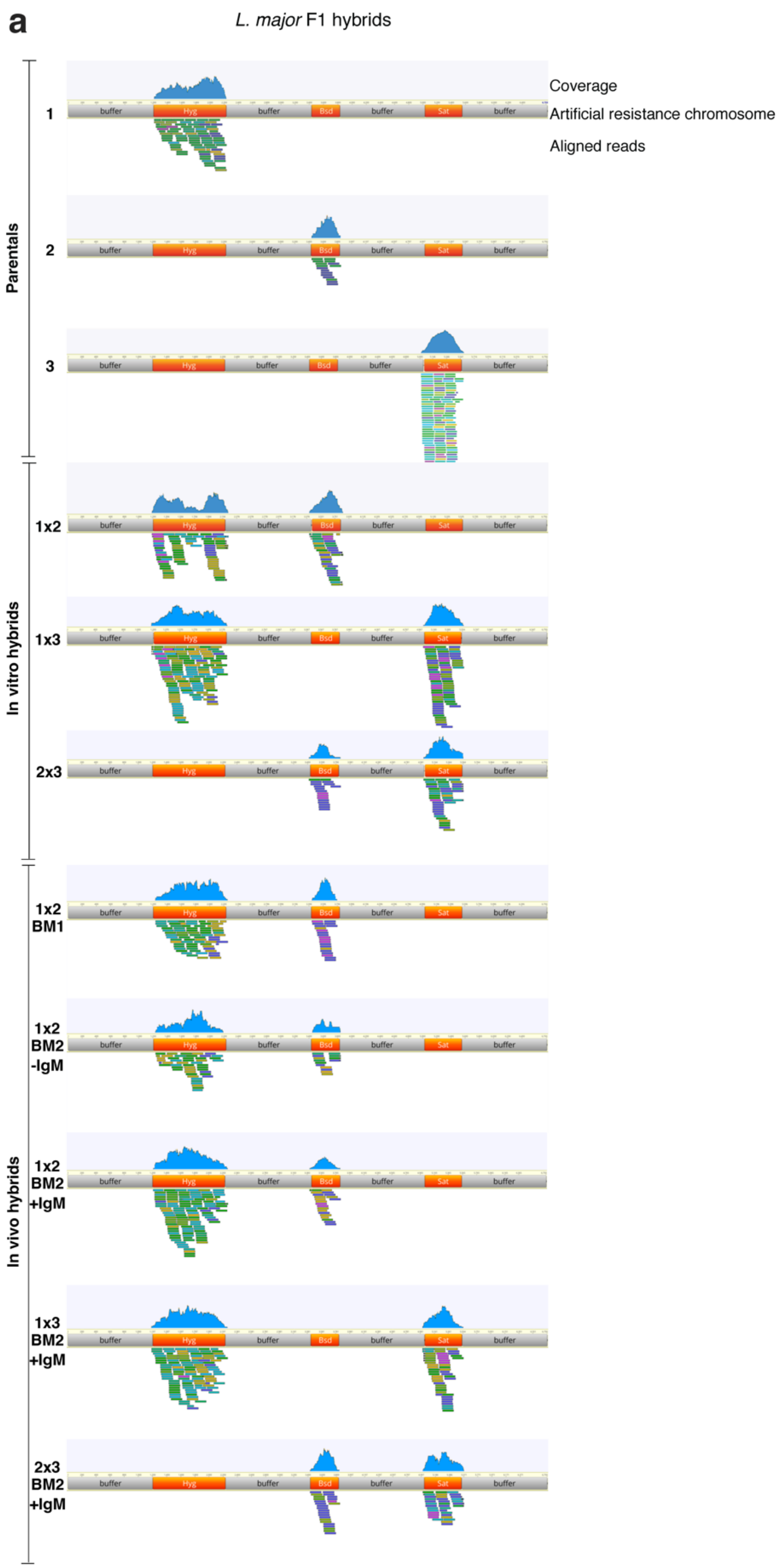

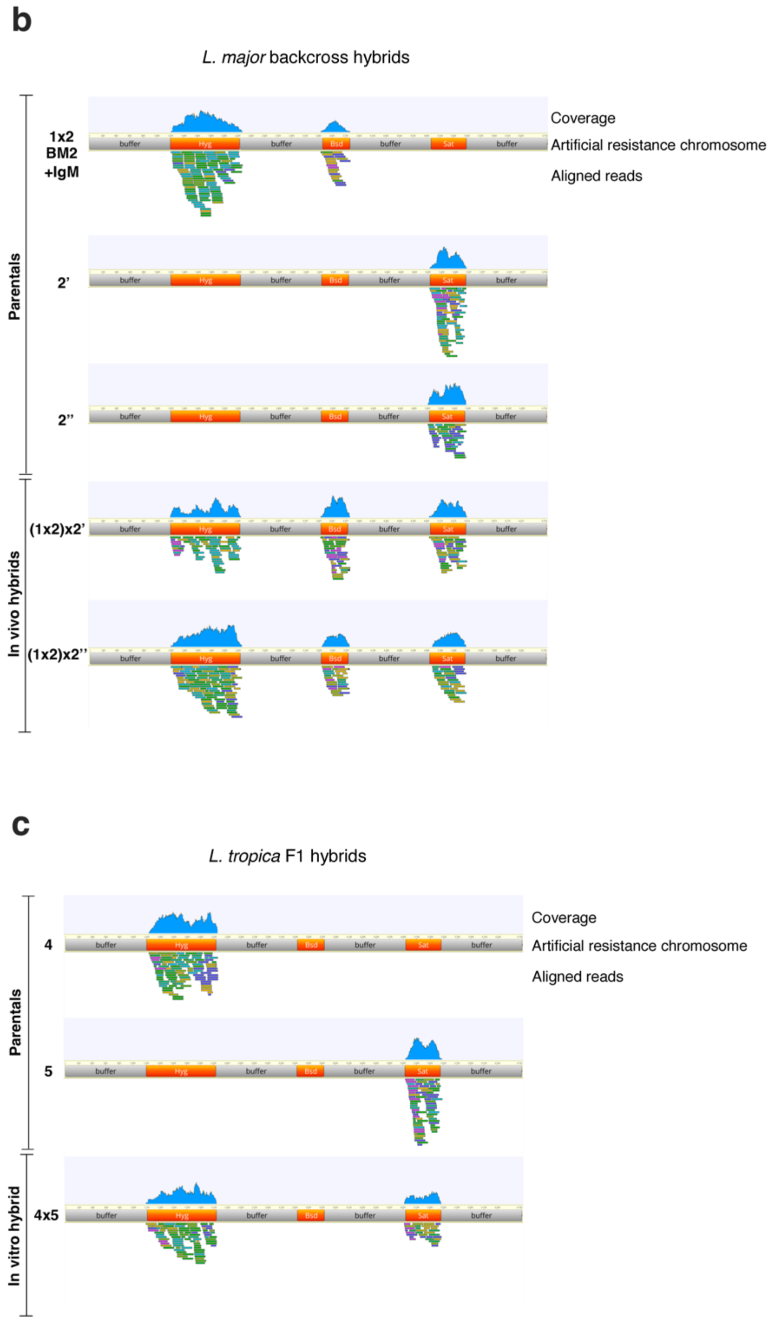


**Supplementary Fig. 8. Resistance markers whole-genome analysis of parental and hybrid lines.** Schematic display of an artificial chromosome containing arbitrary loci of the three antibiotic resistance genes (Hygromycin – HYG, Blasticidin – BSD, Nourseothricin – SAT) used as parental markers in this study. Raw reads were processed and displayed using Geneious prime software v2021.2.2.

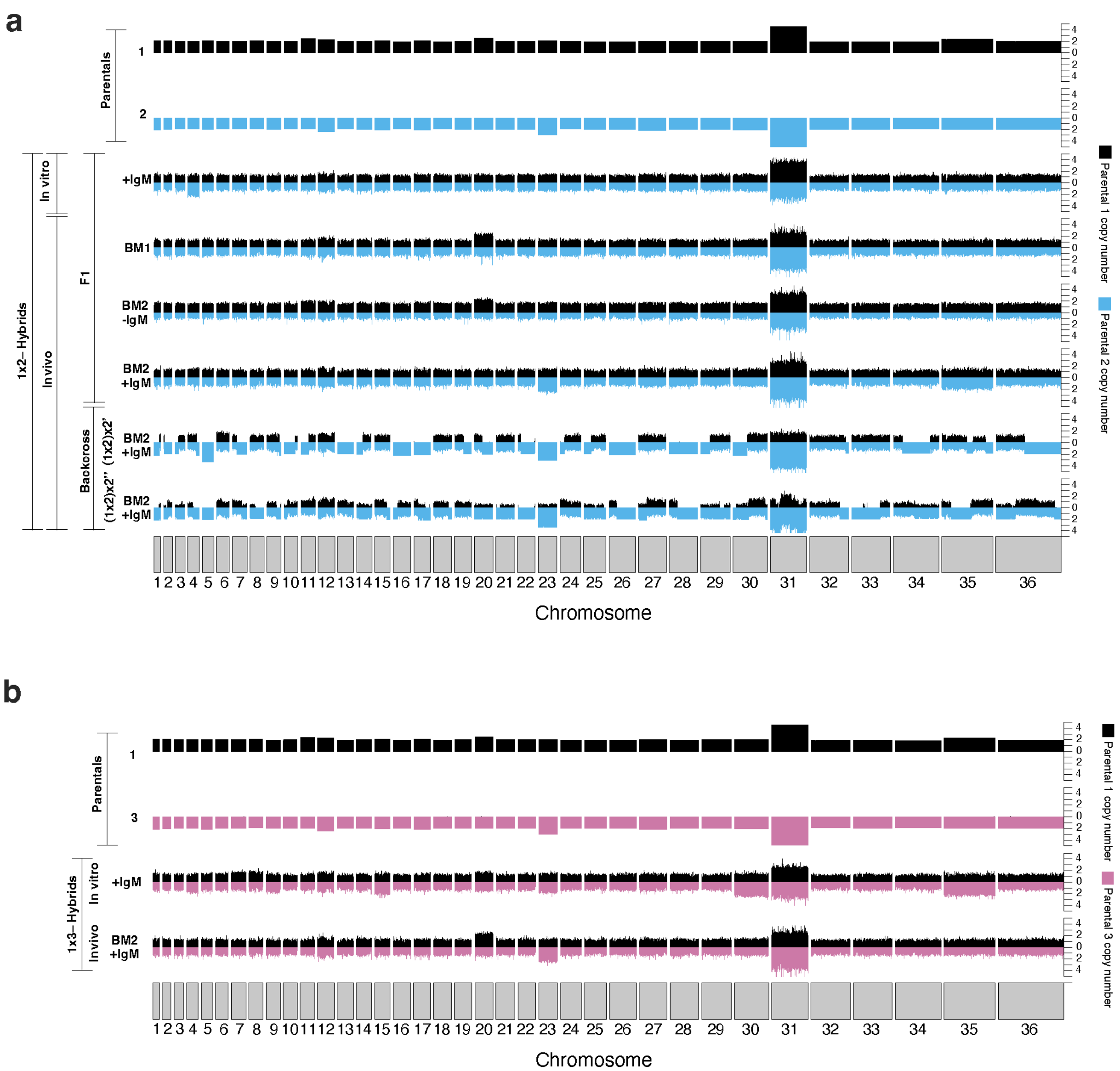


**Supplementary Fig. 9. Whole-genome analysis of *Leishmania major* hybrids. (a, b)** Allele-specific chromosomal copy number as inferred by read depth and allelic proportions. Biparental ancestry was confirmed across the whole genome for crosses between parentals 1x2 **(a)** and 1x3 **(b)**. Biparental inheritance of backcrosses for 1x2 crosses are also shown in Fig. 4d. Parental 1, WR-SSU-HYG; Parental 2, FVI-FKP40-BSD; Parental 3, FVI-FTL-SAT. Parental 2', FV1-FKP40-SAT; Parental 2'', FV1-SSU-SAT.

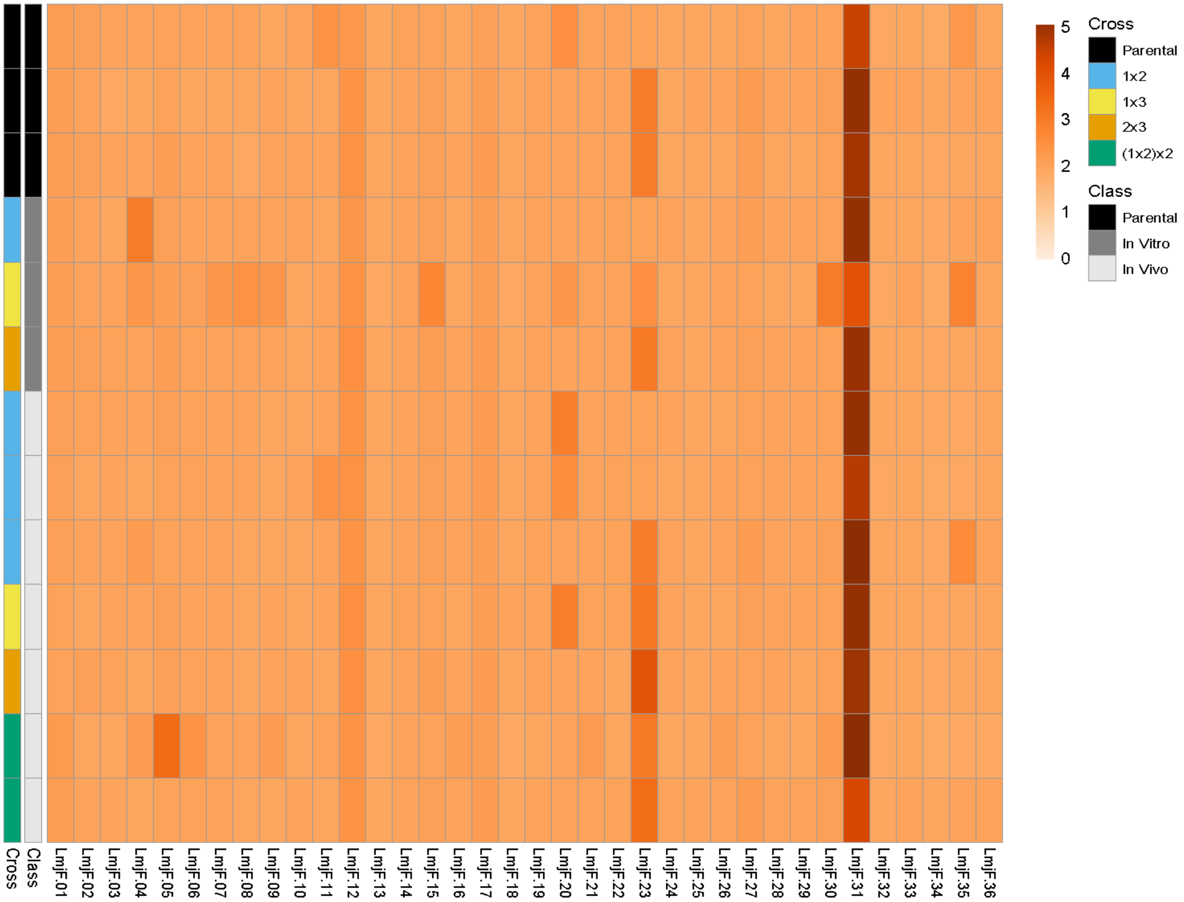


**Supplementary Fig. 10. Somy heatmap of sequenced *Leishmania major* parentals and hybrids.** Whole-genome analysis shows characteristic *Leishmania major* aneuploidy chromosome distribution in all samples. Parental 1, WR-SSU-HYG; Parental 2, FVI-FKP40-BSD; Parental 3, FVI-FTL-SAT. Parental 2', FV1-FKP40-SAT; Parental 2'', FV1-SSU-SAT.

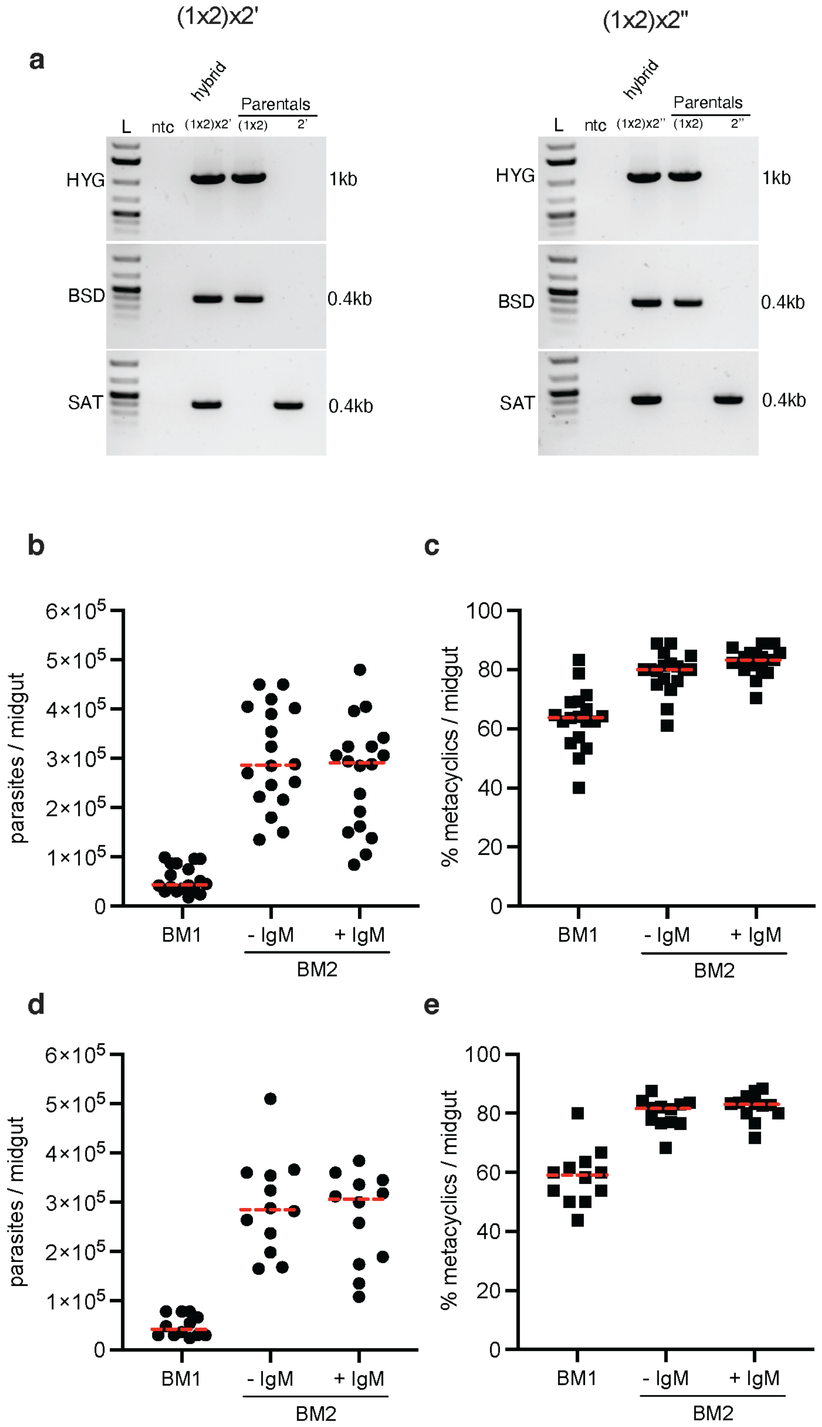


**Supplementary Fig. 11. Hybrid genotyping and infection status in sand flies after a naturally acquired *Leishmania major* infection. (a-e)** Sand flies were fed on mice lesions composed of a parental and a hybrid line**. (a)** *Leishmania major* backcross hybrid genotyping by PCR targeting parental selectable drug markers HYG (Hygromycin), BSD (Blasticidin) and SAT (Nourseothricin). F1 (1x2) hybrid (a cross between parental 1(WR-SSU-HYG) and parental 2 (FVI-FKP40-BSD); Parental 2’, FV1-FKP40-SAT; Parental 2’’, FV1-SSU-SAT.; ntc, no template control; L, 1kb plus ladder (Invitrogen). Triple drug resistant hybrid lines were cloned before genotyping. Only sand flies given a second bloodmeal containing IgM produced hybrids. A single hybrid representative is shown for positive events from each group. Parasite number (b,d) and percentage of metacyclic promastigotes (c,e) in *L. major*-infected *Lu. longipalpis*. At 6 days post-infection, a proportion of the sand flies were provided a second uninfected blood meal in the absence (-IgM) or presence (+IgM) IgM (500 µg/mL). Infection status of individual sand flies was assessed at 14 days after the first blood meal or 8 days after the second bloodmeal. Parental line combination (1x2)x2’ (b,d); (1x2)x2’ (c,e).

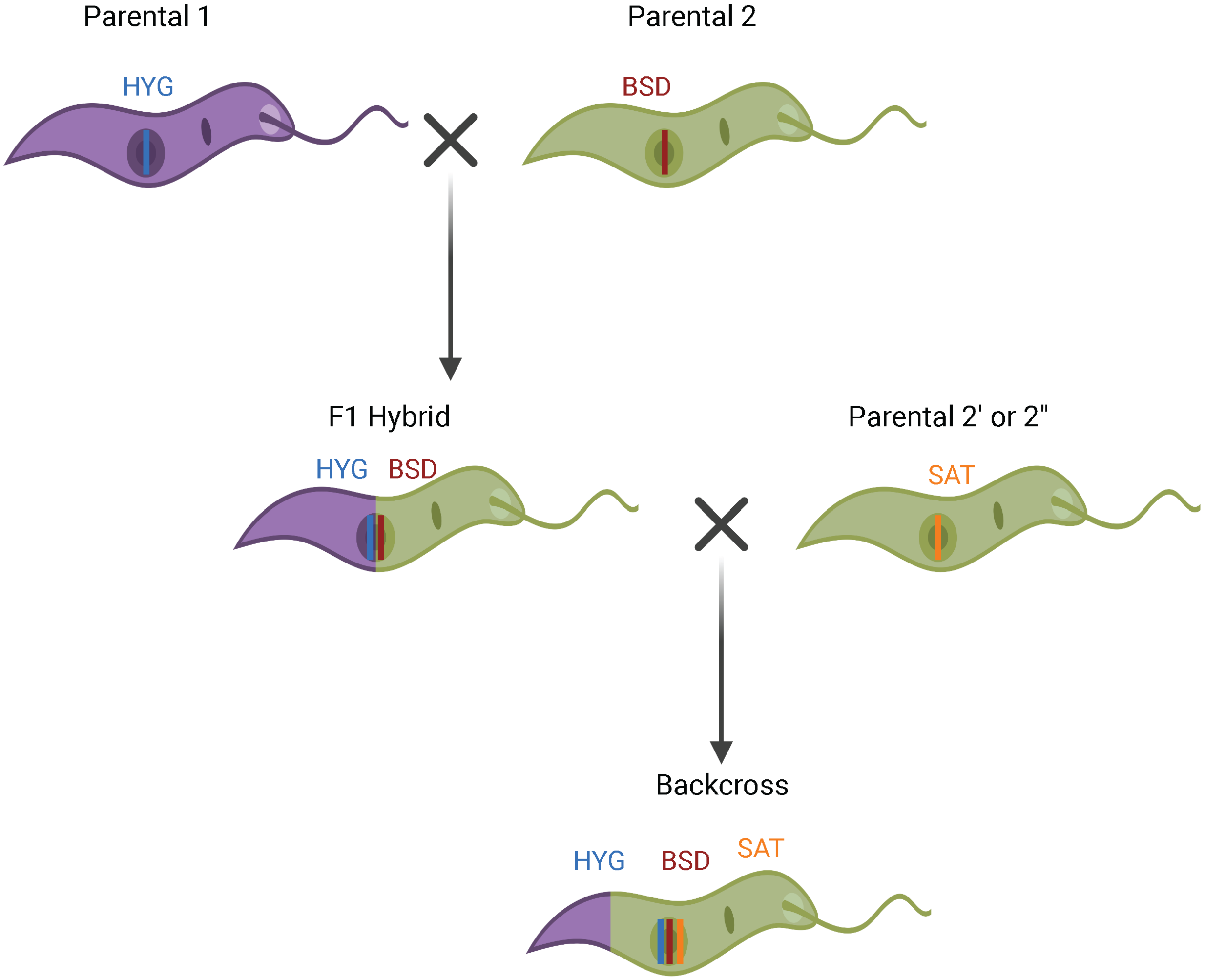


**Supplementary Fig. 12. Diagram outlining our approach to the generation of backcrosses.** *L. major* parental 1 (WR-SSU-HYG) and parental 2 (FVI-FKP40-BSD) resistant to hygromycin (HYG) or blasticidin (BSD), respectively, were crossed to produce F1 hybrids. HYG/BSD double resistant F1 hybrids were backcrossed to *L. major* parental 2' (FV1-FKP40-SAT) or 2" (FV1-SSU-SAT), both resistant to Nourseothricin (SAT) inserted at loci in chromosome 16 and 27, respectively. This resulted in the recovery of fertile backcross hybrids resistant to HYG/BSD/SAT.

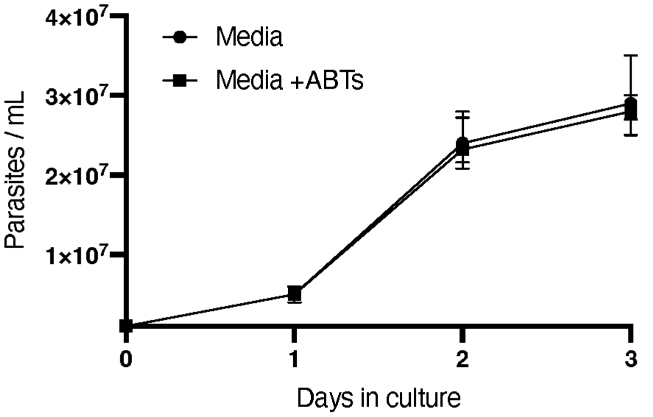


**Supplementary Fig. 13. *L. major* promastigote growth in the presence of an antimicrobial cocktail.** Mid log phase *L. major* promastigotes were cultured in complete Schneider's media in the presence or absence of an antimicrobial cocktail (ABTs) to control the growth of bacteria and fungi during plating of infected sand fly midguts. ABTs: Penicillin-Streptomycin (Gibco, 100 U/mL); Gentamicin (Sigma, 50 µg/mL); Caspofungin (Sigma, 15 µg/mL); 5-fluorocytosine (Sigma, 30 µg/mL).

**Supplementary Table 1. Sand fly blood meal sources.** Literature review of studies that determined the blood source from field-captured fed sand flies.

| Blood meal source | Sand fly *spp.* | References |
|---|---|---|
| **Human** | *Phlebotomus perniciosus* | 76-80 |
| | *Ph. argentipes* | 81,82 |
| | *Ph. orientalis* | 83,84 |
| | *Ph. papatasi* | 85-88 |
| | *Ph. ariasi* | 78 |
| | *Ph. sergenti* | 85,88 |
| | *Lutzomyia longipalpis* | 89-91 |
| **Bovine** | *Ph. perniciosus* | 76,80,88 |
| | *Ph. argentipes* | 81,82 |
| | *Ph. orientalis* | 83 |
| | *Ph. papatasi* | 86-88 |
| | *Ph. sergenti* | 85 |

| | | |
|---|---|---|
| | *Lu. longipalpis* | 91 |
| **Canine** | *Ph. perniciosus* | 77,80 |
| | *Ph. orientalis* | 83,84 |
| | *Ph. papatasi* | 85,86 |
| | *Ph. sergenti* | 85 |
| | *Lu. longipalpis* | 89-92 |
| **Bird** | *Ph. perniciosus* | 76,77,79,80 |
| | *Ph. argentipes* | 81 |
| | *Ph. papatasi* | 87 |
| | *Ph. sergenti* | 85 |
| | *Lu. longipalpis* | 89-92 |
| **Goat** | *Ph. perniciosus* | 76,77 |
| | *Ph. argentipes* | 81,82 |
| | *Ph. orientalis* | 83,84 |
| | *Ph. papatasi* | 88 |
| | *Ph. ariasi* | 77 |
| | *Ph. sergenti* | 85 |
| **Sheep** | *Ph. perniciosus* | 76,78,79,88 |
| | *Ph. orientalis* | 83,84 |
| | *Ph. papatasi* | 88 |
| | *Ph. ariasi* | 77,78 |
| | *Ph. sergenti* | 85 |
| **Horse** | *Ph. perniciosus* | 77-80 |
| | *Ph. ariasi* | 78 |
| | *Ph. sergenti* | 85 |
| | *Lu. longipalpis* | 90 |
| **Rodent** | *Ph. papatasi* | 86-88 |
| | *Ph. sergenti* | 85 |
| | *Lu. longipalpis* | 90 |
| **Cat** | *Ph. perniciosus,* | 77,80 |
| | *Ph. papatasi* | 85 |
| | *Ph. sergenti* | 85 |
| | *Lu. Longipalpis* | 90 |
| **Swine** | *Ph. perniciosus* | 77-80 |
| | *Ph. ariasi* | 78 |
| | *Lu. longipalpis* | 91 |
| **Donkey** | *Ph. perniciosus* | 76,78 |
| | *Ph. orientalis* | 83,84 |
| | *Ph. sergenti* | 85 |
| **Rabbit** | *Ph. perniciosus* | *78-80,93* |
| | *Ph. ariasi* | 78 |
| | *Ph. sergenti* | 88 |
| **Hyrax** | *Ph. papatasi* | 85,87 |
| | *Ph. sergenti* | 85 |

| | | |
|---|---|---|
| **Turkey** | *Ph. perniciosus*<br>*Ph. ariasi* | 76<br>78 |
| **Hare** | *Ph. perniciosus* | 77 |
| **Buffalo** | *Ph. argentipes* | 81,82 |

**Supplementary Table 2. IgM concentration range in normal serum.**

| Animal | IgM - serum range levels (mg/mL) | References |
|---|---|---|
| **Human** | 0.4-2.3 | 94-97 |
| **Bovine** | 1-3 | 98-100 |
| **Mice** | 0.3-0.8 | 96,101 |

**Supplementary Table 3. PCR primers used for genotyping.**

| Target | PCR product (bp) | Forward primer (5'-3') | Reverse primer (5'-3') | Ref. |
|---|---|---|---|---|
| **Hygromycin B (HYG)** | 1026 | GGTAACGGTGCGG GCTGACGCCACCAT GAAAAGCCTGAAC TC | CGAGATCCCACGT AAGGTGCCTATTC CTTTGCCCTCG | 102 |
| **Blasticidin (BSD)** | 393 | ATGCCTTTGTCTCA AGAAGAATC | TTAGCCCTCCCAC ACATAAC | 103 |
| **Nourseothricin (SAT)** | 398 | ACCTATCCGACCAA GGCTTT | CGCTGTTTCGTTC GAGACTT | Current work |

**Supplementary Video 1**. *Leishmania major* metacyclic promastigotes in culture media supplemented with 20% FBS + 5% inactivated adult dog serum. Video taken 30 min after seeding. Event 1. Purified metacyclic promastigotes were used to exclude in vitro multiplication rosettes and for clear visualization of parasite/parasite interaction. Scale bar = 20 µm.

**Supplementary Video 2.** *Leishmania major* metacyclic promastigotes in culture media supplemented with 20% FBS + 5% inactivated adult dog serum. Video taken 30 min after seeding. Event 2. Scale bar = 20 μm.

**Supplementary Video 3.** *Leishmania major* metacyclic promastigotes in culture media supplemented with 20% FBS + 5% inactivated adult dog serum. Video taken 180 min after seeding. Scale bar = 20 μm.

**Supplementary Video 4.** *Leishmania major* stationary phase promastigotes in culture media supplemented with 20% FBS + Bovine IgM (50 μg/mL)**.** Video taken 24 hours after seeding. Scale bar = 50 μm.

**Supplementary Video 5.** *Leishmania major* stationary phase promastigotes in culture media supplemented with 20% FBS + peanut agglutinin (PNA) (50 μg/mL)**.** Video taken 24 hours after seeding. Scale bar = 50 μm.

**Supplementary Video 6.** *Leishmania major* metacyclic promastigotes in culture media supplemented with 20% FBS. Video taken 30 min after seeding. Event 1. Scale bar = 50 μm.

**Supplementary Video 7.** *Leishmania major* stationary phase promastigotes in culture media supplemented with 20% FBS. Video taken 30 min after seeding. Event 2. Scale bar = 50 μm.

**Supplementary Video 8.** In vivo *Leishmania* mating clump formation. Event 1. A sand fly midgut was dissected approximately 30 minutes after imbibing a second uninfected blood meal containing IgM (500 μg/mL)**.** Intact red blood cells are visible. Clumping formation (arrows) follows a pattern similar to that observed in vitro (Supplementary Videos 1,2). Scale bar = 50 μm.

**Supplementary Video 9.** In vivo *Leishmania* mating clump formation. Event 2. A sand fly midgut was dissected approximately 30 minutes after imbibing a second uninfected blood meal containing IgM (500 μg/mL)**.** Intact red blood cells are visible. Clumping formation (arrows) follows a pattern similar to that observed in vitro (Supplementary Videos 1,2). Scale bar = 50 μm.

**Supplementary Video 10.** In vivo *Leishmania* mating clump formation. Event 1. A sand fly midgut was dissected 24 hours after imbibing a second uninfected blood meal containing IgM (500 μg/mL)**.** Partially digested red blood cells are visible. Fully formed *Leishmania* mating clumps (LMCs) can be seen at the center of the video. Scale bar = 50 μm.

**Supplementary Video 11.** In vivo *Leishmania* mating clump formation. Event 2. A sand fly midgut was dissected 24 hours after imbibing a second uninfected blood meal containing IgM (500 μg/mL)**.** Partially digested red blood cells are visible. Fully formed *Leishmania* mating clumps (LMCs) can be seen at the center of the video (arrows). As shown for in vitro LMC formation (Supplementary Videos 1-3), this video captures the fusion of smaller clumps into one large clump. Scale bar = 50 μm.

**Supplementary Video 12.** In vivo *Leishmania* mating clump formation. Event 3. A sand fly midgut was dissected 24 hours after imbibing a second uninfected blood meal containing IgM (500 µg/mL**).** Partially digested red blood cells are visible. Fully formed *Leishmania* mating clumps (LMCs) can be seen at the center of the video (arrows). As showed for in vitro LMC formation (Supplementary Videos 1-3), the fusion of smaller clumps into a bigger one can be observed. Scale bar = 50 µm.

**Supplementary sequence 1.** Genbank sequence file of artificial resistance chromosome used as mapping template on supplementary fig. 8.

**Supplementary references**


76 Remadi, L. *et al.* Molecular detection and identification of Leishmania DNA and blood meal analysis in Phlebotomus (Larroussius) species. *Plos Neglect Trop D* **14**, doi:ARTN e0008077 10.1371/journal.pntd.0008077 (2020).

77 Gonzalez, E. *et al.* Identification of blood meals in field captured sand flies by a PCR-RFLP approach based on cytochrome b gene. *Acta Trop* **152**, 96-102, doi:10.1016/j.actatropica.2015.08.020 (2015).

78 Bravo-Barriga, D. *et al.* Detection of Leishmania DNA and blood meal sources in phlebotomine sand flies (Diptera: Psychodidae) in western of Spain: Update on distribution and risk factors associated. *Acta Trop* **164**, 414-424, doi:10.1016/j.actatropica.2016.10.003 (2016).

79 Maia, C. *et al.* Molecular detection of Leishmania DNA and identification of blood meals in wild caught phlebotomine sand flies (Diptera: Psychodidae) from southern Portugal. *Parasit Vectors* **8**, 173, doi:10.1186/s13071-015-0787-4 (2015).

80 Abbate, J. M. *et al.* Identification of trypanosomatids and blood feeding preferences of phlebotomine sand fly species common in Sicily, Southern Italy. *PloS one* **15**, e0229536, doi:10.1371/journal.pone.0229536 (2020).

81 Garlapati, R. B., Abbasi, I., Warburg, A., Poche, D. & Poche, R. Identification of bloodmeals in wild caught blood fed Phlebotomus argentipes (Diptera: Psychodidae) using cytochrome b PCR and reverse line blotting in Bihar, India. *J Med Entomol* **49**, 515-521, doi:10.1603/me11115 (2012).

82 Poche, D. M. *et al.* Bionomics of Phlebotomus argentipes in villages in Bihar, India with insights into efficacy of IRS-based control measures. *PLoS Negl Trop Dis* **12**, e0006168, doi:10.1371/journal.pntd.0006168 (2018).

83 Aklilu, E. *et al.* Some aspects of entomological determinants of Phlebotomus orientalis in highland and lowland visceral leishmaniasis foci in northwestern Ethiopia. *Plos One* **13**, doi:ARTN e0192844 10.1371/journal.pone.0192844 (2018).

84 Gebresilassie, A. *et al.* Host-feeding preference of Phlebotomus orientalis (Diptera: Psychodidae) in an endemic focus of visceral leishmaniasis in northern Ethiopia. *Parasit Vectors* **8**, 270, doi:10.1186/s13071-015-0883-5 (2015).

85 Valinsky, L., Ettinger, G., Bar-Gal, G. K. & Orshan, L. Molecular Identification of Bloodmeals From Sand Flies and Mosquitoes Collected in Israel. *Journal of Medical Entomology* **51**, 678-685, doi:10.1603/Me13125 (2014).

86 Azizi, K., Askari, M. B., Kalantari, M. & Moemenbellah-Fard, M. D. Molecular detection of Leishmania parasites and host blood meal identification in wild sand flies from a new endemic rural region, south of Iran. *Pathog Glob Health* **110**, 303-309, doi:10.1080/20477724.2016.1253530 (2016).

87 Abbasi, I., Cunio, R. & Warburg, A. Identification of blood meals imbibed by phlebotomine sand flies using cytochrome b PCR and reverse line blotting. *Vector Borne Zoonotic Dis* **9**, 79-86, doi:10.1089/vbz.2008.0064 (2009).

88 Jaouadi, K. *et al.* Blood Meal Analysis of Phlebotomine Sandflies (Diptera: Psychodidae: Phlebotominae) for Leishmania spp. Identification and Vertebrate Blood Origin, Central Tunisia, 2015-2016. *Am J Trop Med Hyg* **98**, 146-149, doi:10.4269/ajtmh.17-0313 (2018).

89 Carvalho, G. M. L. *et al.* Bloodmeal Identification in Field-Collected Sand Flies From Casa Branca, Brazil, Using the Cytochrome b PCR Method. *Journal of Medical Entomology* **54**, 1049-1054, doi:10.1093/jme/tjx051 (2017).

90 Sales, K. G. D. *et al.* Identification of phlebotomine sand fly blood meals by real-time PCR. *Parasite Vector* **8**, doi:ARTN 230 10.1186/s13071-015-0840-3 (2015).

91 Morrison, A. C., Ferro, C. & Tesh, R. B. Host preferences of the sand fly Lutzomyia longipalpis at an endemic focus of American visceral leishmaniasis in Colombia. *Am J Trop Med Hyg* **49**, 68-75, doi:10.4269/ajtmh.1993.49.68 (1993).

92 Sant'Anna, M. R. *et al.* Blood meal identification and parasite detection in laboratory-fed and field-captured Lutzomyia longipalpis by PCR using FTA databasing paper. *Acta Trop* **107**, 230-237, doi:10.1016/j.actatropica.2008.06.003 (2008).

93 Gonzalez, E. *et al.* Identification of blood meals in field captured sand flies by a PCR-RFLP approach based on cytochrome b gene. *Acta Trop* **152**, 96-102, doi:10.1016/j.actatropica.2015.08.020 (2015).

94 Cassidy, J. T. & Nordby, G. L. Human serum immunoglobulin concentrations: prevalence of immunoglobulin deficiencies. *J Allergy Clin Immunol* **55**, 35-48, doi:10.1016/s0091-6749(75)80006-6 (1975).

95 Gonzalez-Quintela, A. *et al.* Serum levels of immunoglobulins (IgG, IgA, IgM) in a general adult population and their relationship with alcohol consumption, smoking and common metabolic abnormalities. *Clin Exp Immunol* **151**, 42-50, doi:10.1111/j.1365-2249.2007.03545.x (2008).

96 Palma, J., Tokarz-Deptula, B., Deptula, J. & Deptula, W. Natural antibodies - facts known and unknown. *Cent Eur J Immunol* **43**, 466-475, doi:10.5114/ceji.2018.81354 (2018).

97 Weitkamp, J.-H., Lewis, D. B. & Levy, O. in *Avery's Diseases of the Newborn (Tenth Edition)* (eds Christine A. Gleason & Sandra E. Juul) 453-481.e457 (Elsevier, 2018).

98 Bayram, B. *et al.* Comparison of immunoglobulin (IgG, IgM) concentrations in calves raised under organic and conventional conditions*. *International Journal of Approximate Reasoning* **50**, 995-999 (2016).
99 Butler, J. E. Bovine Immunoglobulins: A Review. *Journal of Dairy Science* **52**, 1895-1909, doi:https://doi.org/10.3168/jds.S0022-0302(69)86871-2 (1969).
100 Pomorska-Mol, M., Krzysiak, M. K., Larska, M. & Wlodarek, J. The First Report of Immunoglobulin G, M, and A Concentrations in Serum of European Bison and Their Changes with Age. *J Immunol Res* **2020**, 2614317, doi:10.1155/2020/2614317 (2020).
101 Bos, N. A. *et al.* Serum immunoglobulin levels and naturally occurring antibodies against carbohydrate antigens in germ-free BALB/c mice fed chemically defined ultrafiltered diet. *Eur J Immunol* **19**, 2335-2339, doi:10.1002/eji.1830191223 (1989).
102 Akopyants, N. S. *et al.* Demonstration of genetic exchange during cyclical development of Leishmania in the sand fly vector. *Science* **324**, 265-268, doi:10.1126/science.1169464 (2009).
103 Inbar, E. *et al.* The mating competence of geographically diverse Leishmania major strains in their natural and unnatural sand fly vectors. *PLoS Genet* **9**, e1003672, doi:10.1371/journal.pgen.1003672 (2013).
